# Temporal resource variation promotes growth equalization through alternative plasticity strategies

**DOI:** 10.64898/2026.08.22.746401

**Authors:** David M. Ekkers, Marina Costa Rillo, Stefany Moreno-Gámez, Oscar P. Kuipers, G. Sander van Doorn

**Author notes:** Corresponding author. David Matthias Ekkers, Waardenburg Ecology, Oosterweg 127, 9751 PE, Haren (Groningen), The Netherlands.

## Abstract

Evolutionary theory predicts that fluctuating environments favor adaptations that maximize geometric-mean fitness by reducing variance in performance across conditions. We tested this prediction experimentally by evolving the lactic acid bacterium Lactococcus cremoris on the sugars fructose and galactose in density-controlled chemostats under four resource regimes: constant supply of fructose, constant supply of galactose, a constant mixture of both sugars, and a temporally alternating supply of the two. In the absence of temporal variation, trade-offs between fructose and galactose resulted in evolutionary divergence into a fructose specialist and a galactose specialist. In contrast, adaptation to temporal resource variation equalised growth performance on both sugars by increasing its growth rate on galactose and decreasing it on fructose. Interestingly, performance equalization emerged across replicate populations through distinct resource transition strategies indicated by differences in resource affinity and growth recovery on fructose and galactose among strains isolated from the evolved populations. Our results show that temporal resource variation selects for variance-minimizing resource adaptations while adopting multiple resource transition strategies, illustrating how distinct modes of metabolic plasticity can yield convergent fitness outcomes.

## 1. Introduction

### 1.1 The theoretical framework of evolutionary responses to temporal variation

Organisms inhabiting temporally varying environments face the challenge of maintaining a steady energy supply despite fluctuations in the type and availability of resources. Unlike spatially varying environments, where organisms can exploit geographical refuges from unfavorable conditions, temporally varying environments inevitably impose periods of low fitness (Gottschal et al., 1979; Reboud and Bell, 1997; Kassen, 2002). To persist, organisms must evolve traits that minimize variance in fitness across conditions while simultaneously improving average performance. Strategies that achieve high mean performance but perform poorly under one environmental state are unlikely to persist long-term, as they risk extinction during adverse periods (Bulmer, 1994). Consequently, in fluctuating environments, evolutionary fitness is determined by performance across all environmental states and, under suitable assumptions, can be approximated by the geometric mean of fitness (Gillespie, 1973). Thus, when the potential for generic adaptations that increase overall performance has been exploited, individuals maximize fitness by equalizing performance across environments (Bradshaw, 1965).

Phenotypic strategies that enable survival in temporally varying environments generally fall into three categories: (1) generalists that exhibit a fixed trait representing an optimal compromise across conditions; (2) phenotypically plastic strategies that express alternative traits in response to environmental cues; and (3) bet-hedging strategies that stochastically express traits optimized for different environmental states, thereby ensuring reliable transmission of the genotype by at least a subset of offspring (a risk-spreading tactic). At the level of the genotype, each of these strategies confers a higher long-term geometric mean fitness than a specialist optimized for a single environmental condition. While such specialists may achieve the highest arithmetic mean fitness across environmental states, their geometric mean fitness is typically lower due to their high variance in performance (Futuyma and Moreno, 1988; Gavrilets and Scheiner, 1993; Leroi et al., 1994; Reboud and Bell, 1997; Kassen, 2002; Buckling et al., 2007; Turner and Elena, 2000). Which of these alternative strategies evolves depends on the nature of the trade-offs governing performance across fluctuating conditions (Fig. 1a,b). If trade-offs are weak, evolution may favour a generalist with a fixed phenotype that performs sufficiently well across all conditions (Kassen and Bell, 1998). In contrast, strong trade-offs preclude viable compromise strategies and instead induce phenotypic polymorphisms. In predictable environments, polymorphisms can arise through plastic switching, where organisms adjust the expression of traits in response to informative environmental cues. By contrast, in unpredictable and/or highly variable environments, bet-hedging strategies may be favoured based on stochastically switching phenotypes (Veening et al., 2008). Lastly, negative frequency-dependence and mild fluctuations provide favourable conditions for the maintenance of a genetic polymorphism of two specialists that express different, complementary phenotypes. These alternative outcomes are associated with different patterns of maximum growth rate, variance in growth rate, and switching costs across conditions (Fig. 1c). A generalist has low temporal variance in growth rate and no switching costs, but also a relatively low maximum performance on any specific resource. A phenotypically plastic strategy reduces temporal variance in growth rate and achieves moderate-to-high performance across resources, but is likely to incur costs of switching between strategies (Murren et al., 2015) or maintenance costs for the ability to sense and respond to environmental cues. A polymorphism of two specialists is expected to achieve the highest growth performance on each resource without switching costs. However, each specialist exhibits high variance in performance and is, therefore, more susceptible to population bottlenecks or extinction during adverse periods. Importantly, the efficacy of plastic strategies depends not only on how much plasticity can be expressed (reaction-norm magnitude) but also on how fast it can be expressed (Dupont et al., 2024). These temporal features of plasticity, which shape the efficiency of plastic resource transitions, are expected to affect geometric-mean fitness in fluctuating environments.

**Figure 1.**
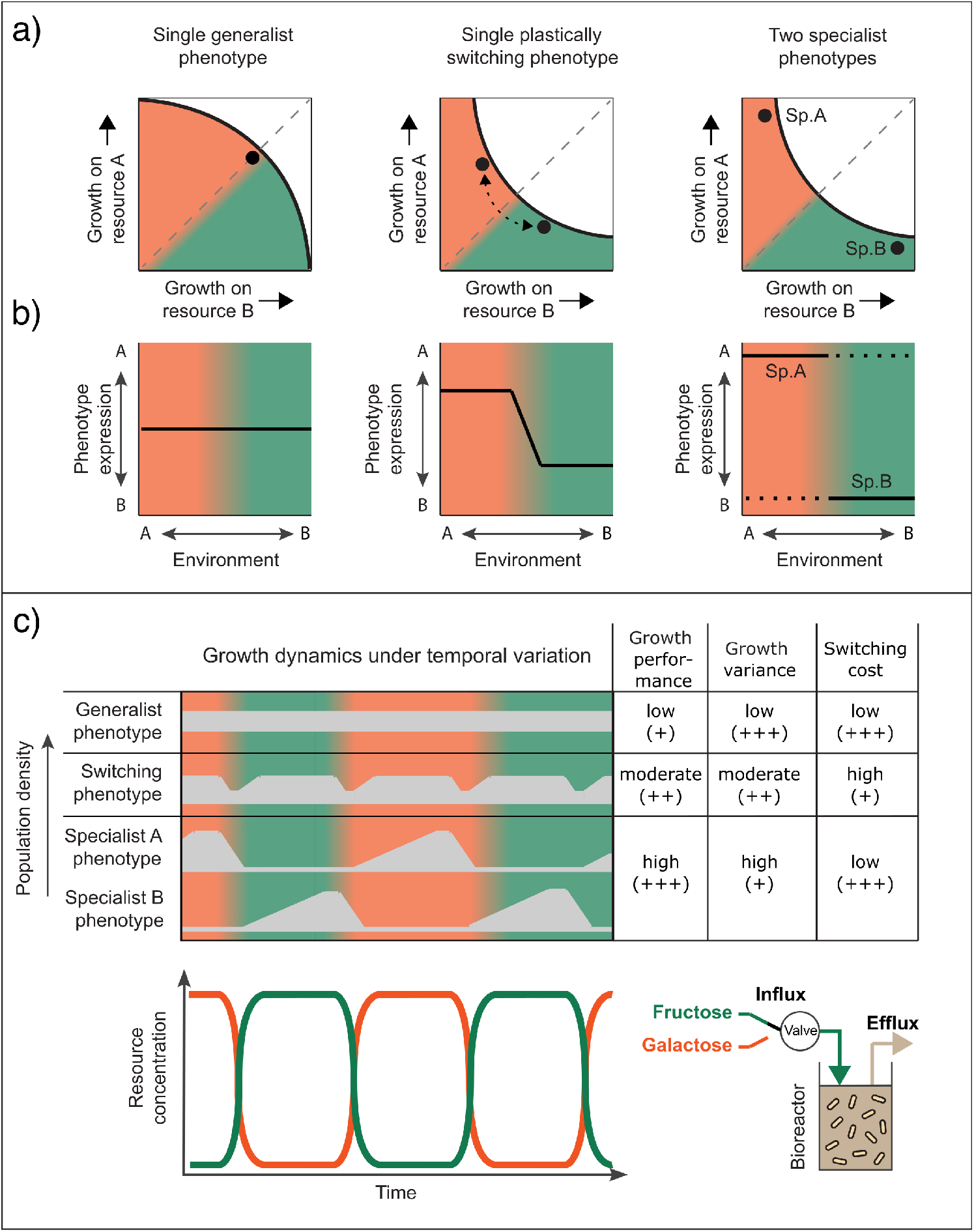
**a)** Three alternative phenotypic strategies under a two-resource regime of temporal variation. Colored shaded areas indicate regions in phenotype space adapted to growth on resource A (orange) or resource B (green). Black lines indicate trade-off curves delineating the boundaries between evolutionary feasible (colored) and unfeasible (white) phenotypes. Black dots indicate the realised phenotype. The grey dashed diagonals mark the region in phenotype space where growth performance on the two resources is equal. “Sp.” refers to specialist genotypes. **b)** The reaction norm of the three phenotype strategies in response to changes in resource availability. The dashed line of the right panel indicates poor growth performance. **c)** Upper panel: Predicted growth dynamics of the phenotypic strategies in response to temporally alternating resource availability with impact on different growth performance aspects (max growth, growth variance, and switch cost), plusses indicate the level of positive fitness effects of different growth characteristics. Bottom panel displays a schematic visualisation of the continuous culture selection regime temporal variation in two resources.

### 1.2 Evolution experiments are constrained by a limited understanding of trade-offs and selection pressure

Efforts to experimentally investigate the mechanisms governing adaptation to temporal variation have yielded mixed results. Studies have reported the evolution of generalists in response to temporal variation (Condon et al., 2014; Duncan et al., 2011; Ketola et al., 2013), cost-free generalists (Buckling et al., 2007; Reboud and Bell, 1997; Kassen and Bell, 1998), single specialists (Cooper and Lenski, 2010), a polymorphism of multiple specialists (Gottschal et al., 1981) or the co-evolution of specialists and generalist phenotypes (Legan et al., 1987). This apparent inconsistency among experimental evolution studies on temporal variation suggests that the evolutionary outcome of temporal variation depends on the strength of trade-offs among traits under selection. Trade-offs in resource utilisation strategies dictate the costs of switching between resources and the variance in growth performance (Fig. 1c) and therefore influence the evolutionary outcome of populations in temporally varying environments. Thus, our ability to interpret the evolution of different phenotype-determination strategies under temporal variation ultimately relies on our understanding how these trade-offs interact with the temporal structure of selection. Such a selection structure may include resource transition speed, costs of plasticity, speed of plasticity, and strength of trade-offs-all factors are expected to affect variance in fitness across cycles. In this context, the selection regime considered under a finer temporal timescale can be divided into two phases: (1) A steady state phase where growth improvement on separately encountered resources at specific concentrations is selected for; (2) a resource transition phase where resource concentrations change dynamically in time, favoring continuous expression adjustment (plasticity) to optimize growth performance, minimize growth variance and switching cost (Fig. 1c). From a mechanistic perspective, it remains unclear to what extent improvements in fitness components during the steady-state and resource-transition phases trade off with one another, and how these trade-offs occur.

In the context of microbial evolution experiments, mixed outcomes under temporally varying environments may partly reflect the widespread use of the serial-transfer culturing method (Reboud and Bell, 1997; Ketola et al., 2013; Buckling et al., 2007; Gottschal et al., 1981; Duncan et al., 2011; Kassen and Bell, 1998). Its simplicity, cost-effectiveness, reproducibility, and reliability make it the generally preferred culturing method for experimental evolution. However, an important disadvantage of serial dilution is its inherently temporally variable selection regime (Gresham and Dunham, 2014; Ekkers et al., 2020). Cultures are diluted daily and grown to the stationary phase, so that organisms experience cyclical changes in nutrient availability (from high to low), shifts in pH, and accumulation of metabolic waste products. Together, these fluctuating conditions expose organisms to a multidimensional selection regime that periodically selects for fast metabolic startup, high growth rate, high growth yield, resistance to pH changes, waste-product stress resistance, and growth recovery. This multidimensional regime in serial-transfer culturing makes it difficult to disentangle the relationship between induced selection and the resulting adaptations. By contrast, continuous cultures present a solution, providing a steady-state selection regime that minimises spatio-temporal variation (Ekkers et al., 2020). Moreover, systematically varying continuous-culturing conditions over time allows for the precise induction of defined regimes of temporal variation, which can then be compared to control treatments without temporal variation (i.e., standard continuous cultures). Such a design allows probing the mechanistic linkage between selection regime, evolved phenotypes, and genetic changes.

Here, we use continuous-culture evolution experiments to investigate how metabolic trade-offs shape adaptation to temporally varying environments. We evolved the bacterium *Lactococcus cremoris* for more than 500 generations in a regime of temporal resource variation induced by an alternating supply of fructose and galactose (Fig. 1d). Temporal variation was accurately induced under continuous culture conditions by modulating resource supply rates, while maintaining a constant temperature, pH, and average population density throughout the experiment. Three steady-state control treatments were used to compare the evolutionary outcomes of the temporally varying treatment: two in which cultures were supplied with only one sugar (either fructose or galactose), and one in which cultures were supplied with a constant mixture of both sugars. Imposing selection on fructose and galactose metabolism provides a suitable model system to probe the constraints of adaptation to temporal variation, because efficient growth on these two sugars is subject to a strong trade-off in *L. cremoris*: fructose and galactose impose conflicting demands on glycolytic flux optimisation (Ekkers et al., 2022). Thus, we expect that, under a temporally varying environment, the metabolic trade-off between fructose and galactose will prevent the evolution of a cost-free generalist, leading instead to divergent adaptations in the utilisation of each sugar. We also expect an improved ability to switch between metabolic states optimised for each resource. By measuring growth rates, their variance, and phenotypic plasticity of the evolved strains, including time-resolved switching and concentration-dependent reaction norms, we reconstructed the evolutionary trajectories of the populations and revealed hidden variation in molecular mechanisms and reaction norms underlying phenotype determination.

## 2. Materials and Methods

### 2.1 Bioreactor system

The evolution experiment was performed using a custom-built bioreactor system (Ekkers et al., 2020), consisting of 100 ml glass reactor vessels (*Duran 100 ml GL45*) with a stainless steel headplate. The bioreactors are kept in a waterbath (*Julabo MB*) with submersible magnetic stirrers (*Cimarec micro VWR 442-4530*) for a controlled, constant temperature and constant mixing of the culture. The pH in the bioreactors is regulated by a computer-controlled PID loop using a pH sensor (*Applisens pH sensor for mini bioreactor 8 mm*) and a peristaltic pump (*Ismatec REGLO ICC MS-4/8*), which adds base (35% NaOH) to the culture. In- and out-flux of (spent) media is controlled by high accuracy peristaltic pumps (*Ismatec IPCN-24*). Temporal variation in sugar availability was induced by a set of computer-controlled pinch valves (*Sirai 3/2 NC-NO solenoid pinch valve*). The whole system is controlled using a computer (*NI CompactRIO*) with a custom LabView graphical interface, which controls and logs: flux rate, pH, temporal variation, and temperature.

### 2.2 Experimental procedures of the evolution experiment

Because long-term continuous culture evolution experiments are vulnerable to biofilm formation, we avoid commonly used model organisms such as *Escherichia coli* or *Bacillus subtilis*, and instead chose *Lactococcus cremoris* MG1363 (formerly *L. lactis* subsp. *cremoris*, Li et al. 2021), which has a very low propensity to form biofilms. Cultures were grown in 60 ml chemically-defined medium for prolonged cultivation (CDMPC) (Price et al., 2019). In the temporal treatment, the bioreactors were supplemented through time with either 0.5% (wt/v) fructose (F-CDMPC) or 1% (wt/v) galactose (G-CDMPC) as a source of carbon and energy. The mix treatment was supplemented with a mix of 0.25% wt/v fructose + 0.5% wt/v galactose, the only-fructose treatment with 0.5% fructose, and the only-galactose treatment with 1% wt/v galactose. The 1:2 fructose:galactose wt/v ratio was chosen because of the high growth-rate asymmetry between fructose and galactose for the ancestral *L. lactis* strain, which is pre-adapted for efficient growth on glucose (Ekkers et al., 2022). The twofold higher supply of galactose served to partially balance performance on both sugars at the start of the experiment, and to avoid a large initial asymmetry in fitness between conditions that would endanger survival of the culture in the temporal treatment. Cultivation was performed anaerobically (25 ml/min N2 headspace gasflow) at a pH of 6.5 under continuous dilution and stirring (330 rpm) at 30°C. The starting dilution rate was set at 0.2 for the temporal and only-galactose treatment and 0.3 for the mix and only-fructose treatment. These differences between starting dilution were also based on the asymmetry between the ancestral performance on fructose and galactose (Ekkers et al., 2022). Bioreactor dilution rates were monitored daily, and gradually increased over the course of the experiment to control culture density (0.5 ¿OD_600_ ¡1.0) and maintain consistent selection for higher growth rate. Every day, 1.5 ml samples were drawn from each bioreactor to make −80°C glycerol stocks. Weekly samples were taken to screen for infections using flow cytometry and growth plate analysis with glucose-supplemented M17-medium agar plates.

Temporal variation was induced in the bioreactors by alternating the influx of fructose- and galactose-supplemented medium. Throughout the experiment, the temporal variation interval was set at three generation times (3/the dilution rate of the bioreactors). The control treatments were induced by continuously pumping media supplemented with a mix of both fructose and galactose (mix treatment), only fructose (only-fructose treatment) or only galactose (only-galactose treatment). Each treatment was run with four parallel independent replicates (Fig. 2a). Total duration of the treatments were: temporal 665, fructose 1111, galactose 565 and mix 575 generations. This unequal number of generations was due to growth rate differences among treatments. We sampled populations of each replicate of each treatment across three timepoints (T1, T2, T3), which were selected per treatment, in order to be equally spread over the growth rate performance increase achieved throughout each treatment.

**Figure 2.**
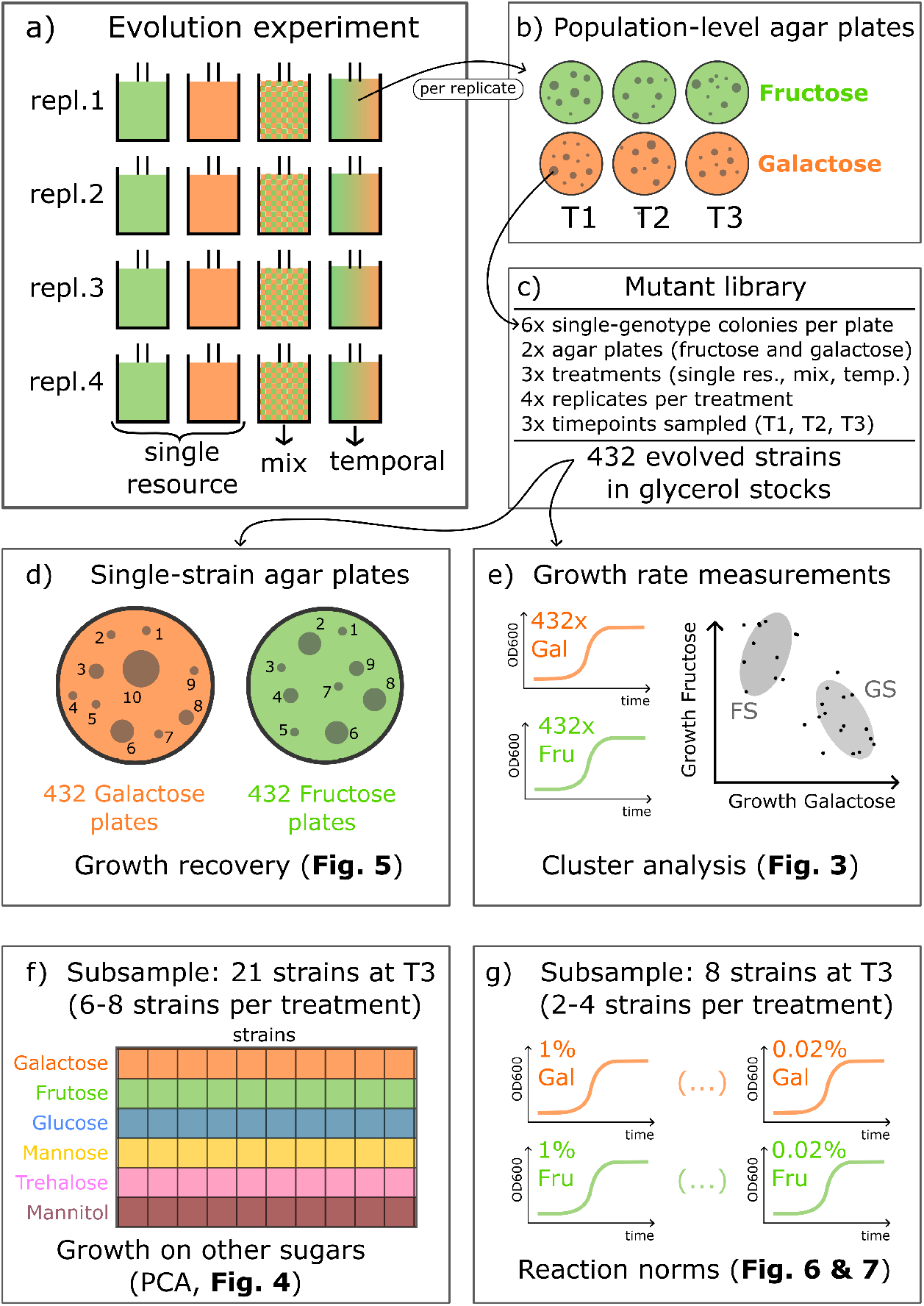
Overview of the methodology. Results of analyses are referenced by figure numbers. **a)** Twenty-four chemostats were assigned to three treatments (four replicates each): *single resource* - two separated chemostats, one with galactose (orange) and one with fructose (green); *mix* - both sugars present simultaneously; *temporal* - single sugar environments alternating between galactose and fructose every three generations. **b)** Population-level samples were taken from each chemostat at three timepoints along the experiment (namely T1, T2, and T3) and plated on both resources: fructose and galactose. This procedure resulted in 3 treatments x 4 replicates x 3 timepoints x 2 agar plates = 72 agar plates (CDMPC). **c)** From each of these agar plates, 6 colonies were selected, resulting in a mutant library of 432 strains. **d)** Single-strain plate analysis was conducted by plating each strain on each sugar and quantifying CFU count and size; **e)** cluster analysis; GS galactose specialist, FS fructose specialist; **f)** subsample T3: 21 strains; growth on other sugars; **g)** subsample T3: reaction norms.

### 2.3 Analysing pH sensor data

Bacteria acidify their culture medium as they break down sugars. In *L. cremoris*, the rate of acidification is proportional to metabolic activity and growth, so we used pH decline as a proxy for population-level metabolism. The pH of each chemostat was maintained near constant by drip-feeding NaOH via a peristaltic pump that responded to decreases in pH. Although acidity remained close to pH 6.5, high-resolution sensor data revealed a fine-scale sawtooth pattern (Supp. Fig. S7), reflecting rapid increases in pH from NaOH addition alternating with gradual acidification due to bacterial metabolism. Because pH declined linearly between NaOH additions, we estimated metabolic activity by fitting a linear least-squares regression to each interval (Supp. Fig. S6). The gradual decrease in pH reflects the buildup of acidic waste metabolites that triggered NaOH supply, whereas the sudden increase marks the addition of a NaOH droplet. Thus, both the frequency of NaOH addition and the slope of acidification serve as proxies for growth rate. Measurements were taken at the end of each cycle to avoid interference from metabolic shifts. This analysis primarily establishes relative differences between fructose and galactose metabolism at a given time point (Supp. Fig. S8). Comparisons across longer timescales must be made cautiously, as adjustments of the dilution rate alter the buffering capacity of the medium.

### 2.4 Mutant library construction and phenotypic analysis on agar plates

After the evolution experiment was completed, a library of evolved strains was created to phenotypically characterise the evolved mutants in each replicate of each treatment across the three timepoints (T1, T2, T3) (Fig. 2b,c). Population samples from the bioreactors were taken from the glycerol stock, diluted 500 times; 50 *µ*l of the diluted cell suspension was then plated on F-CDMPC (fructose 1% wt/v) and/or G-CDMPC (galactose 1% wt/v) agar plates. Population samples from the only-fructose treatment were plated on F-CDMPC plates, from the only galactose treatment on G-CMDPC plates, and from the temporal and mix treatments on both types of plate (Fig. 2b). After 48 hours of incubation at 30°C. We picked 48 colonies per treatment and timepoint to construct a library of single genotypes from the population samples (i.e., twelve genotypes were sampled from each replicate population (six colonies were picked from both the F-CDMPC and G-CDMPC) from the temporal, mix, and single resource treatments (six genotypes from the fructose patch and six from the galactose patch per timepoint). To sample the population-sample plates as broadly as possible for all occurring phenotypes, we chose to pick the six colonies in a way to include contrasting sizes from each plate: two large, two average-sized and two small colonies. Each colony was then grown separately overnight at 30°C without shaking in 2ml F-CDMPS or G-CDMPC (depending on the treatment it derived from), and glycerol stocks were prepared from these cultures, yielding a library of 432 evolved strains (Fig. 2c).

### 2.5 Growth recovery experiments

The single-genotype glycerol stocks from the mutant library were inoculated F-CDMPC plates for the only-fructose treatment, G-CDMPC for the only-galactose, and on both resources (F-CDMPC and G-CDMPC) for the temporal and mix treatments. This re plating step enabled us to quantify the performance of each genotype on both sugars and to monitor the effect of preculturing on growth recovery in the temporal and mix treatments. After 48h of incubation, plates were photographed (*Canon MVX100i*) on a colony counter. The photographs from single-genotype plates were analyzed with the OpenCFU software to measure total colony count and size (area pixel count) for each individual colony (Fig. 2d). To quantify the relative performance of the evolved single-genotypes on fructose and galactose, we calculated the ratio between the total colony count on the F-CDMPC plate and on both plates combined (F-CDMPC + G-CDMPC) (Fig. 5); this (F / (F + G)) ratio was also computed for the median colony size (Supp. Fig. S5).

### 2.6 Growth curves for single genotypes and phenotypic clustering analysis

Growth rates on fructose and galactose were measured for all genotypes from the mutant library after pre-culturing them overnight in F-CDMPC and G-CDMPC. The −80°C stocks were diluted 100 times in PBS, 1 *µ*l of this cell suspension was used to inoculate 100 *µ*l of fresh fructose, galactose and mixed sugar CDMCP (pH=6.5). Each strain was grown in triplo in 384-wells plates (*Greiner Bio-one 781906*) under anaerobic conditions (*VIEWseal Greiner Bio-one*) at 30°C in a plate reader (*Tecan F200*). Estimates of the instantaneous growth rate were obtained by performing local linear regression analyses on the growth curve data, using a sliding window of five data points (measurements were taken every 10 min). After eliminating noisy data from the initial growth phase (OD ¡ 0.16), we determined the maximum growth rate from the regression curves. The three maximum growth rate values were then averaged for each genotype on each sugar.

We used a model-based clustering method to identify and classify the different phenotypic groups that evolved during the evolution experiment (Fig. 2e). The maximum growth rate of every single genotype was plotted as a two-dimensional coordinate: maximum growth rate on fructose on the y-axis and maximum growth rate on galactose on the x-axis, per timepoint per replicate per treatment. To estimate the number of clusters (phenotypic groups), we applied normal (Gaussian) mixture models and a maximum likelihood approach using the R package *mclust* (version 5.4, Scrucca et al. 2016). To consistently apply the same clustering model to all the treatments and timepoints, we selected the EVV model, which allows clusters of ellipsoidal shape (i.e., bivariate Gaussian distributions) with different covariance structures (i.e., different orientations) (Supp. Fig. S2). This EVV model was either the best model selected by the Bayesian Information Criterion (BIC, maximum likelihood corrected for model complexity) or yielded the same number of clusters as the best model selected by BIC independently for each timepoint and treatment. We limited the maximum number of clusters to three, corresponding to the phenotypic groups: fructose specialist, galactose specialist, and generalist.

### 2.7 Growth on a variety of sugars (principal component analysis)

A set of 21 strains was selected from each treatment at timepoint T3: eight genotypes from the temporal, six from the mix, four from the only-galactose, and three from the only-fructose. This selection includes at least three representative strains from each phenotypic cluster (FS, GS) from each treatment. The strains were grown in triplo in batch on fructose, glucose, mannose, galactose, trehalose, or mannitol supplemented (1% wt/v) CDMPC (Fig. 2f), as previously described for fructose and galactose. We performed a principal component analysis (PCA) on the growth values after subtracting the ancestral growth rate from the corresponding evolved strain growth values and scaling the variables to unit variance. PC1 explained 62.4% of the variation in the data, while PC2 explained 20.5%.

### 2.8 Diauxic shift experiments

The mutant strains were precultured in 150 *µ*l of 0.2% wt/v F-CDMPC or G-CDMPC. When the growth curve approached the end of the exponential phase (OD between 0.20 and 0.25) culture was supplemented with 50 *µ*l of CDMPC supplemented with 2% wt/v fructose or galactose. All other culturing conditions were identical to those used for the growth curve analysis performed on the mutant library.

### 2.9 Reaction norm experiments

All culturing conditions were identical to those used for the growth curve analysis performed on the mutant library, except for the inclusion of a wider range of sugar concentrations (0.02, 0.04, 0.07, 0.1, 0.2, 0.5 and 1% wt/v) in sugar-supplemented CDMPC (Fig. 2g).

## 3. Results

### 3.1 Chemostat sensor data confirms growth performance improvements of evolved strains and reveals preferential switching

To assess growth improvements during the evolution experiment, we analyzed culture acidification rates from pH-sensor data in the temporal treatment chemostats. The acidification rate reflects the excretion of metabolic waste products (principally lactate, acetate, and formate) resulting from growth and, thus, serves as a proxy for population-level (i.e., chemostat-level) growth rate. Linear regression analysis of pH sensor measurements revealed a consistent increase in acidification rates across all replicates in the temporal treatment (Supp. Fig. S6), indicating continuous adaptive improvements throughout the experiment. Analysis of the acidification rate of the temporal treatment cultures shows that *L. cremonis* initially has a higher acidification rate on fructose than on galactose (G:F ratio 1:2.7; Supp Fig. S7), consistent with the ancestral pre-adaption to glucose (Ekkers et al., 2022). However, this growth asymmetry reverses in the second half of the experiment in all replicates (G:F ratio 1.8:1; Supp Fig. S7), suggesting that the evolving populations improve their performance on galactose relatively more than on fructose.

### 3.2 Phenotypes from the temporal treatment converge toward equal growth performance

The reconstructed evolutionary trajectories of the mutant library (Fig. 2e) show the emergence of two main phenotypic groups: fructose (FS) and galactose (GS) specialists (Fig. 3). In all four replicates of the mix treatment (i.e., constant availability of both resources), a polymorphism of FS and GS evolved, with FS being more abundant than GS at the end of the experiment (timepoint T3, Fig. 3b). The trajectories of the only-fructose treatment were shorter (i.e., remained closer to the ancestor), less consistent, yet all populations converged towards FS, showing a mild improvement in fructose performance relative to the ancestor (Fig. 3c). This modest improvement likely reflects the ancestral strain’s pre-adaptation to glucose, which is, from a metabolic-pathway perspective, more similar to fructose than to galactose (Ekkers et al., 2022). The temporal (Fig. 3a) and the only-galactose (Fig. 3d) treatments follow similar evolutionary trajectories. At T1, both diverged into a polymorphism: one phenotype showed improvement on galactose coupled with a strong trade-off on fructose, whereas the other improved on both sugars (only-galactose) or primarily on fructose while slightly compromising galactose growth (temporal treatment). At T2, however, only the phenotypes that traded off fructose performance remained in both treatments. The consistency of this pattern across all replicates suggests that adaptation to galactose is constrained, with further improvement only possible at the cost of fructose performance, consistent with previously established metabolic trade-offs between fructose and galactose (Ekkers et al., 2022). At T3, both treatments showed further improvements on galactose, accompanied, interestingly, by simultaneous improvements on fructose. This equalisation of performance on both sugars, absent in the mix treatment (Fig. 3b), occurred in all four replicates of the temporal treatment and in three of four replicates of the only-galactose treatment (Fig. 3a,d). Notably, one of these three replicates of the only-galactose treatment evolved a high-performance generalist phenotype with high growth rate on galactose and fructose (0.9h^−1^ in both sugars at T3; Fig. 3d), despite fructose growth not being under selection in this treatment. These results suggest that the strength of performance trade-offs in galactose specialists weakens as the populations evolve. Nonetheless, the strong consistency of evolutionary trajectories across replicates and treatments suggests that trade-offs constrained alternative evolutionary routes to equalized growth that did not initially involve a decrease in fructose performance.

**Figure 3.**
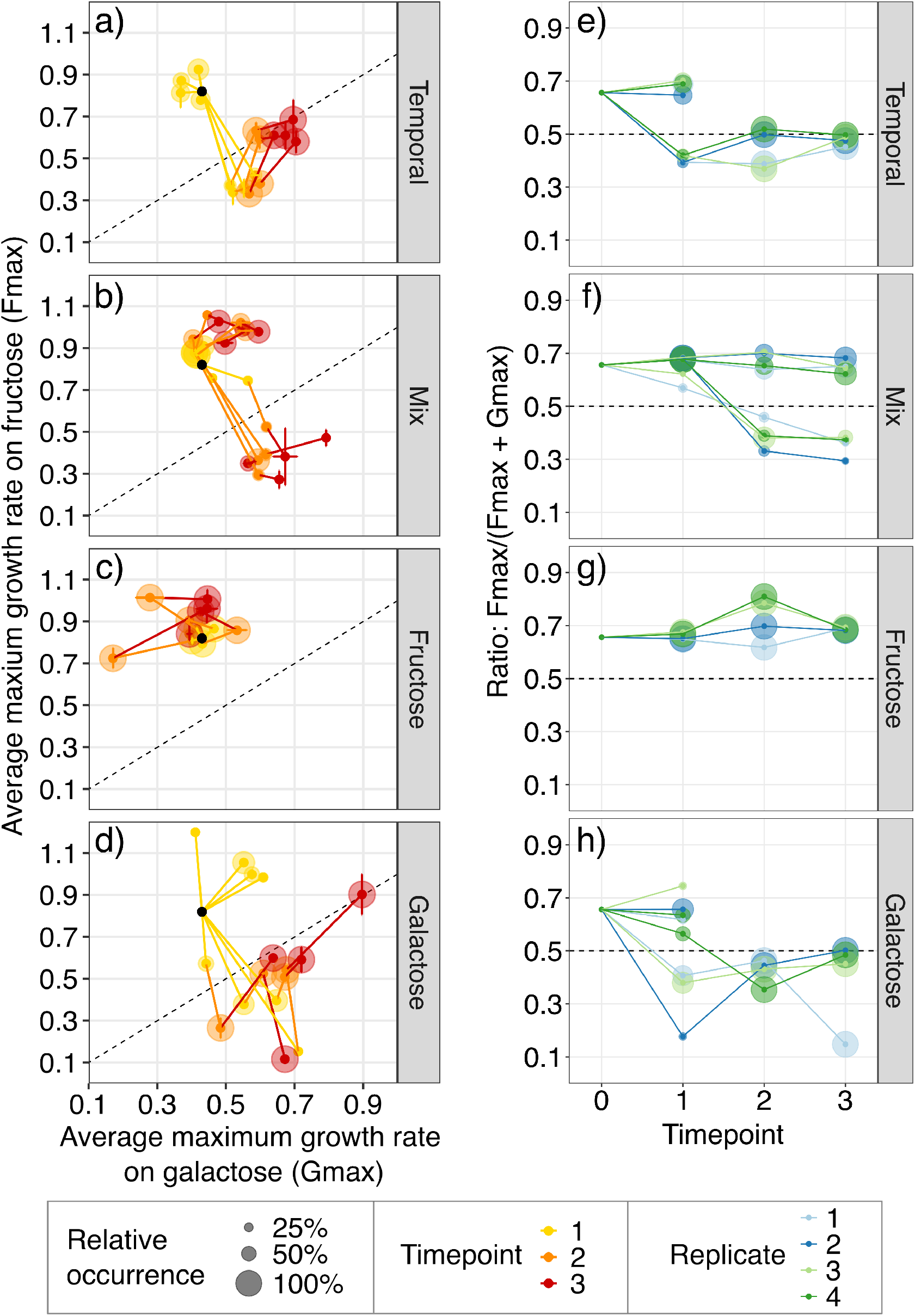
Evolutionary phenotypic trajectories of the four replicates of each treatment quantified by the average maximum growth rates on fructose (Fmn) and galactose (Gmn) **(panels a-d)**, and the ratio between maximum growth rates on fructose and galactose **(panels e-h).** Treatments: temporal (a, e), mix (b, f), only fructose (c, g), and only galactose (d, h). The black dot indicates the ancestral strain. Coloured dots show the median growth values of each phenotypic cluster within each replicate, and error bars show their standard deviations (in both dimensions). Circle sizes represent the relative frequency of each phenotypic cluster within each replicate. (a-d) The grey dashed lines indicates growth-rate equalisation between fructose and galactose, delineating three strategies: fructose specialist (left upper corner), galactose specialist (right bottom corner), and generalists (right top corner and left bottom corner, i.e., extremes of the dashed line).

### 3.3 Phenotypes that evolved in the temporal treatment resemble galactose specialists

To further investigate whether the evolved populations in the temporal treatment resemble galactose specialists from the control treatments, we quantified, for 21 evolved strains at T3 (Fig. 2f), their growth rate on six sugars - fructose, galactose, glucose, mannose, trehalose, and mannitol - relative to the ancestor (Fig. 4). The adaptation to fructose or galactose shows a clear pleiotropy with the other sugars: improvement on fructose leads to improvement on glucose and mannose, whereas improvement on galactose leads to improvement on mannitol and trehalose. Fructose and galactose specialists lie at opposite ends of the primary axis of variation (PC1, Fig. 4). Strains from the mix treatment fall between these two extremes - consistent with the polymorphisms observed in this treatment - while strains from the temporal treatment cluster closer to the galactose specialists.

**Figure 4.**
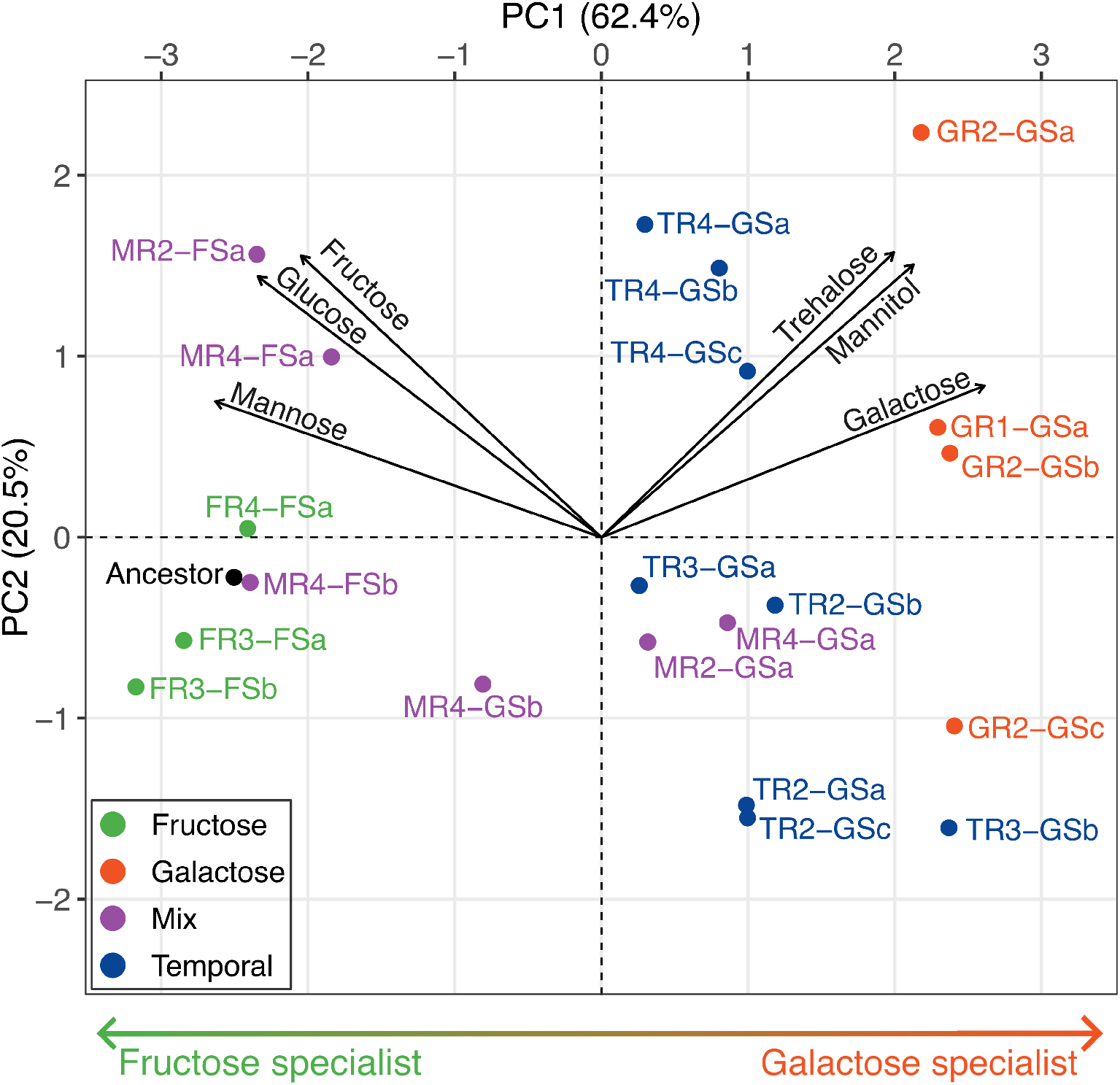
Metabolic characterisation of 21 evolved strains. The first principal component (PC1) explains 62.4% of the variation in growth rates in six sugars (fructose, glucose, mannose, galactose, trehalose, and mannitol; see Methods). PC1 reflects the degree of specialisation towards fructose (green arrow below) and galactose (orange arrow below). Strain labels follow the scheme: the first letter indicates the treatment (M = mix; F = only-fructose; G = only-galactose; T = temporal), followed by the replicate number (R1-R4). FS or GS after the dash denotes fructose or galactose specialists, respectively, according to the cluster analysis (Fig. 3a-d), and the final lowercase letter identifies the strain. Black dot indicates the ancestor strain, which resides in the glucose, mannose fructose side of PC1, consistent with its preadaptive state to glucose. The 21 strains were sampled at the final timepoint (T3), see Methods and Fig. 2f.

### 3.4 Temporal treatment populations evolve alternative plastic strategies

To investigate growth plasticity across replicates of all treatments, we quantified growth recovery - defined as the ability of single cells to resume growth on a given resource - by plating all 432 strains of the mutant library on fructose and galactose media and counting and measuring the resulting colonies (Fig. 2d and Methods). Growth recovery showed considerable variation among the evolved replicate populations in the temporal treatment (Fig. 5; colony size showed consistent results: Supp. Fig. S5). Temporal-treatment replicates 1 (TR1) and 2 (TR2) showed higher growth recovery on galactose than on fructose, particularly when strains were pre-cultured on galactose - patterns similar to the galactose specialists from the only-galactose treatment. In contrast, strains pre-cultured on fructose exhibited greater strain-specific growth recovery, with an average growth recovery between strains that was more equal between sugars. Temporal-treatment replicate 3 (TR3) exhibited a dimorphism linked to the pre-culturing conditions (Fig. 5, timepoint T3): one phenotype showed high growth recovery on fructose, and the other on galactose. This TR3 dimorphism was also evident in the PCA (Fig. 4) as indicated by the separation of strains TR3-GSa (high growth recovery on fructose) and TR3-GSb (high growth recovery on galactose). Interestingly, the TR3 high growth recovery on fructose phenotype (Fig. 5) was more associated with the galactose (GS) than the fructose specialists (FS) in the other analyses (Fig. 3, 4). Lastly, temporal-treatment replicate 4 (TR4) showed a distinct pattern of phenotypes with similar growth recovery on both sugars (Fig. 5, timepoints T2 and T3) and similar growth rate on other non-selected sugars associated with a consistently weaker trade-off on fructose, glucose, and mannose (Fig. 4). Thus, TR4 strains show a more equalised performance across resources, being equally able to start up on both sugars independent of pre-culture conditions (Fig. 5), while the other replicates of the temporal treatment show more dependency on pre-culturing conditions and evolve galactose specialism (TR1, TR2) or a dimorphism of galactose or fructose specialism (TR3).

**Figure 5.**
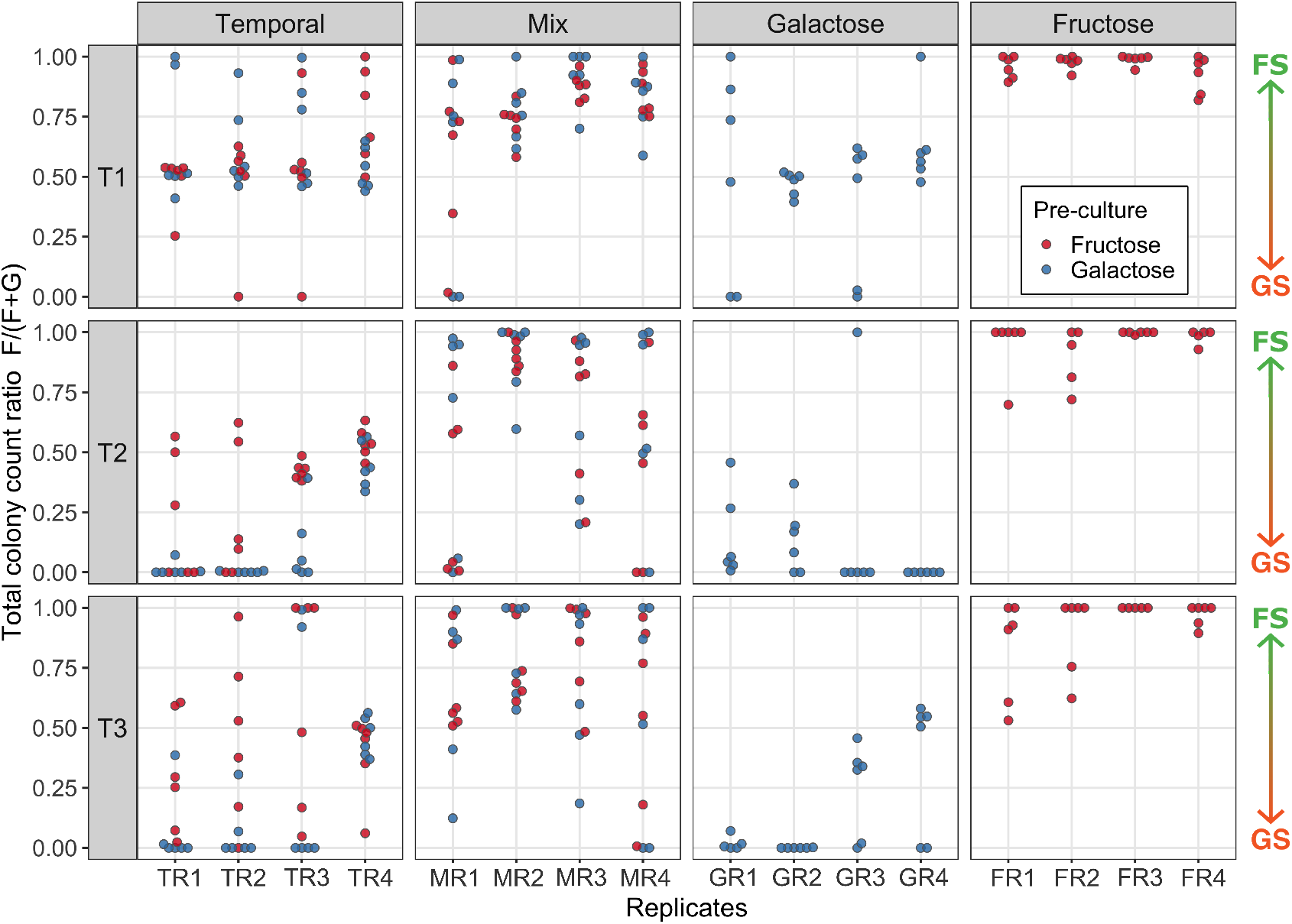
Relative growth recovery of 432 evolved strains on fructose- and galactose-supplemented medium. Colony forming units (CFU) count on fructose was divided by total count to create a ratio that indicates relative growth recovery between both resources (1 = only able to resume growth on fructose, 0 = only able to resume growth on galactose, and 0.5 = equal ability to resume growth on both sugars). Populations are labeled according to the following convention: the first letter indicates the treatment (M = mix; F = only-fructose; G = only-galactose; T = temporal), followed by the replicate number (R1-R4). The green-orange arrows on the right side of the panels indicate the resource specialisation gradient between fructose (FS) and galactose (GS) specialists. T1, T2 and T3 indicate the three sampled timepoints.

The single resource-control treatments showed a clear and expected trait divergence in the growth recovery essays: fructose specialists are more viable on fructose, and galactose specialists are more viable on galactose. The mix-control treatment shows divergence patterns, but weaker than the galactose-only and fructose-only controls. Particularly, the growth recovery trade-off in the galactose specialist is less evident and seems to retain relatively high growth recovery on fructose. Although differences between replicate populations in the mix and single resource control treatments were detected, these differences were generally more consistent among replicates within the same treatment and reflected the specialist groups observed in the evolutionary trajectories of these treatments (Fig. 3).

In conclusion, the temporal treatment displays the highest variability in growth recovery among replicates (Fig. 5), despite showing the most consistent growth-rate evolutionary trajectories among replicates (Fig. 3). Thus, although the temporal selection regime selects for converging growth performance among replicates presumably (Fig. 3), it provides greater scope for alternative plastic strategies to evolve (Fig. 4, 5). This apparent convergence-divergence contrast between growth performance and plasticity could reflect adaptation to the steady state (static) versus resource transition (dynamic) selection regimes experienced under temporal resource variation.

### 3.5 Improvements and trade-offs in resource utilization efficiency evolve at distinct concentration ranges

Given the variation in resource-dependent plasticity (Fig. 5), we asked whether these patterns relate to selection during the resource-transition phase. We reason that the ability to maintain or resume growth across different resource concentrations could influence how effectively strains switch between resources. To test this, we quantified phenotypic change across a gradient of resource concentrations (i.e., each strain’s reaction norm) by measuring expressed growth rates (Fig. 2g, 6, Supp. Fig. S4). These measurements on resource utilization efficiency serve as an approximation resource affinity under Monod kinetics. One representative strain was selected from each phenotypic group identified in Fig. 5 for each treatment.

The measured reaction norms broadly reflect the patterns observed in the growth rate trajectories (Fig. 3) and growth recovery assays (Fig. 5). Surprisingly, resource utilization efficiency changed at specific concentration ranges rather than uniformly across the resource gradient (Fig. 6). For strains with growth rate increases on fructose (four strains: TR3-GSa, TR4-GSa, FR3-FSa and MR4-FSa), the largest relative growth rate increase consistently evolved at low (0.1%) fructose concentration, while weaker growth rate increases (FR3-FSa and MR4-FSa) or even declines (TR3-GSa, TR4-GSa) evolved at higher resource concentrations. Strains with increased growth rates on galactose (all seven measured strains except FR3-FSa) display growth rate increases at higher (0.2-1%) resource concentrations that are spread out more evenly across the resource concentration gradient compared to fructose. Four strains (TR2-GSb, TR3-GSa, TR3-GSb, MR4-GSb) displayed both increases (0.2-1%) and declines (0.07-0.1%) in growth rate within the galactose resource gradient.

**Figure 6.**
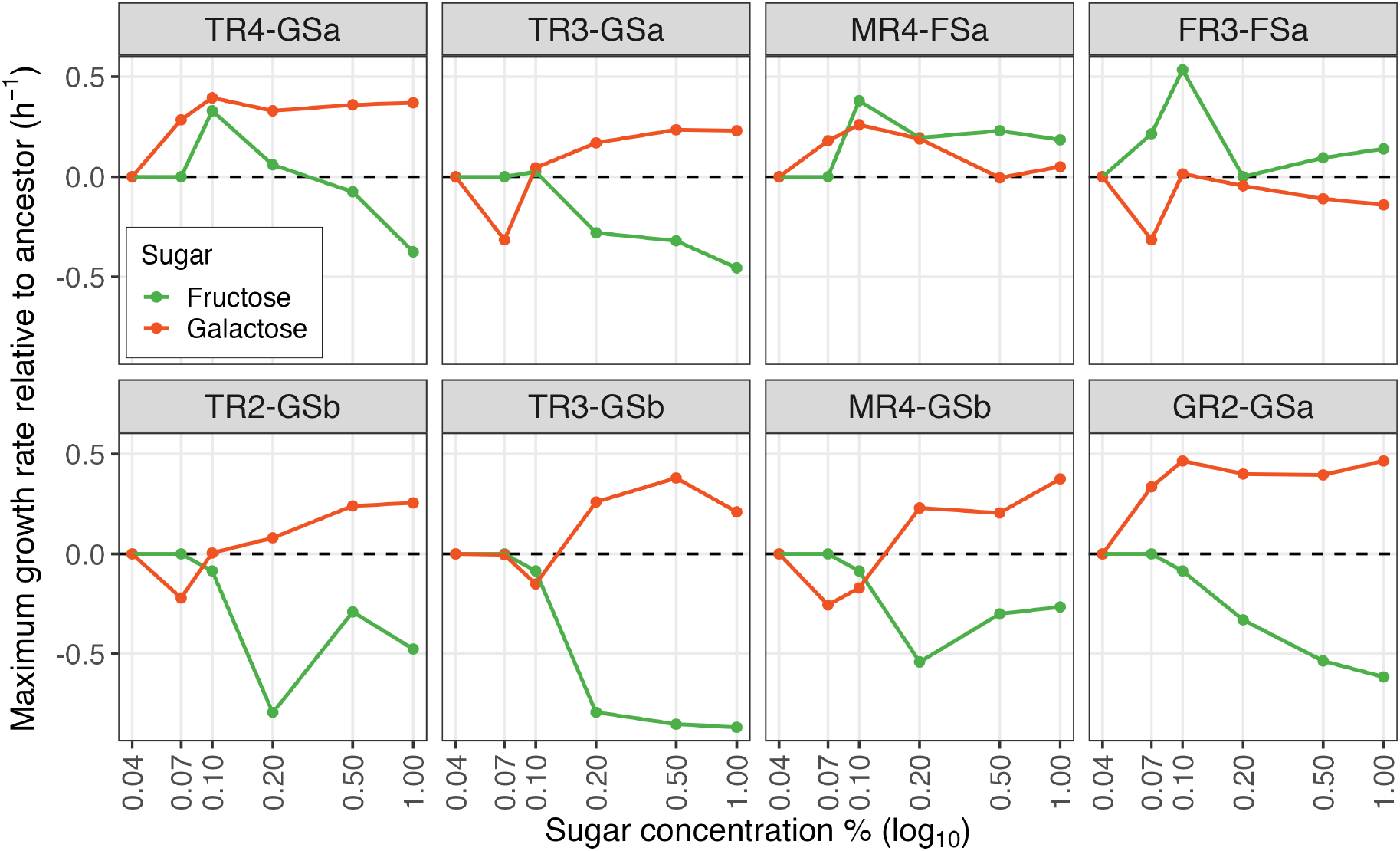
Changes in resource utilization efficiency compared to the ancestor. The maximum rate of growth was measured at a gradient of resource concentrations (% w/v) of the ancestor and eight evolved strains in CDMPC. Ancestral growth rates at each concentration were subtracted from the evolved strains to indicate positive or negative growth changes along the resource gradient. Strain labels follow this scheme: the first letter indicates the treatment (M = mix; F = only fructose; G = only galactose; T = temporal), followed by the replicate number (R1-R4). FS or GS after the dash denotes fructose or galactose specialists, respectively, and a final lowercase letter identifies the strain. Strains were sampled at the final timepoint (T3). No growth was measured in all strains at 0.04% sugar. See Supp. Fig. S3 for relative fold-change in growth rate compared to ancestor, and Supp. Fig. S4 for absolute values.

In TR4-GSa, we observe a distinct pattern: the strain shows a substantial increase in growth rate at low resource concentrations for both carbon sources. This pattern contrasts with the other strains from the temporal treatment, which only exhibit increased growth on galactose and only at higher concentrations. These differences suggest that TR4-GSa may differ from the other temporal strains in its resource-switching behavior.

### 3.6 Metabolic transitions between resources depend both on growth persistence and switch speed

To investigate whether the plastically switching phenotypes in the temporal replicates evolved different plastic switching behaviour, we compared the resource switching efficiency of strains from replicates TR2 and TR4 during transitions between the two sugars. We chose representative strains from these two replicates because they consist of a single resource specialist phenotype but show distinctly different growth recovery and resource utilization efficiency patterns (Fig. 5). TR4 is particularly interesting because its population displayed a ‘generalist’ phenotype in terms of growth recovery on fructose and galactose as well as resource utilization efficiency improvements at relatively low concentration ranges for both sugars (Fig. 5, timepoints T2 and T3), suggesting faster switching between sugars, leading to a potential advantage in temporally varying environments. TR2, in contrast, evolved a galactose specialist phenotype that displays a higher growth recovery on galactose and growth affinity improvements only galactose at higher resource concentrations (Fig. 5), and is expected to show a growth lag when transitioning between sugar strategies.

To investigate resource transition behaviour of TR2 and TR4, we assessed acidification rates immediately following the switch in resource supply (Supp. Fig. S8). Both populations exhibit a temporary reduction in metabolic rate following a switch from fructose to galactose, but no such effect was detected in the reverse transition from galactose to fructose, consistent with other replicates (Supp. Fig. S8). This pattern indicates that resource transition time depends on the direction of the sugar transition. Populations switching from fructose to galactose exhibit a notable retardation in metabolic activity, which can either be explained by a heterogeneous population with different switching strategies or by a homogeneous, plastically-switching population that experiences reduced growth while re-directing metabolic investment toward galactose metabolism.

Single-strain culture analysis of TR2-GSb and TR4-GSa shows a diauxic shift during the transition from fructose to galactose (Fig. 7a), indicating that the resource transition results in a reduction in net growth performance. A fast switch to galactose (max. growth rate on galactose ±1.5 hour after galactose addition) was measured for TR2-GSb culture with a relatively low growth rate and yield on galactose (Fig. 7a). The TR4-GSa culture, in contrast, displayed a delayed shift (max. growth rate on galactose ±2.5 hours after galactose addition) combined with a high growth rate and yield on galactose (Fig. 7a). Whether the observed differences in resource transition time between both clonal populations resulted in synchronized plasticity or are related to heterogeneity switching ability within the clonal cultures (Solopova et al., 2014) remains unclear. Superficially, the resource transition speed measured in the growth curves shows a different pattern from that observed in the resource transition interval during the chemostat acidification curves (Supp. Fig. S8), where we find that TR2 has a longer resource transition interval than TR4. However, based on the delayed diauxic shift and high metabolic yield of TR4-GSa (Fig. 7a), we hypothesize TR4-GSa can switch later, resulting from an ability to metabolise fructose at lower concentrations. Comparing the resource utilization efficiency for TR2-GSb and TR4-GSa on fructose and galactose across a range of sugar concentrations indicates that TR4-GSa consistently sustained a higher growth rate at low concentrations of both sugars compared to TR2-GSb (Fig. 7b). During a resource switch, this means that TR2-GSb is expected to be forced to switch at an earlier stage (at a higher residual sugar concentration) than TR4-GSa.

**Figure 7.**
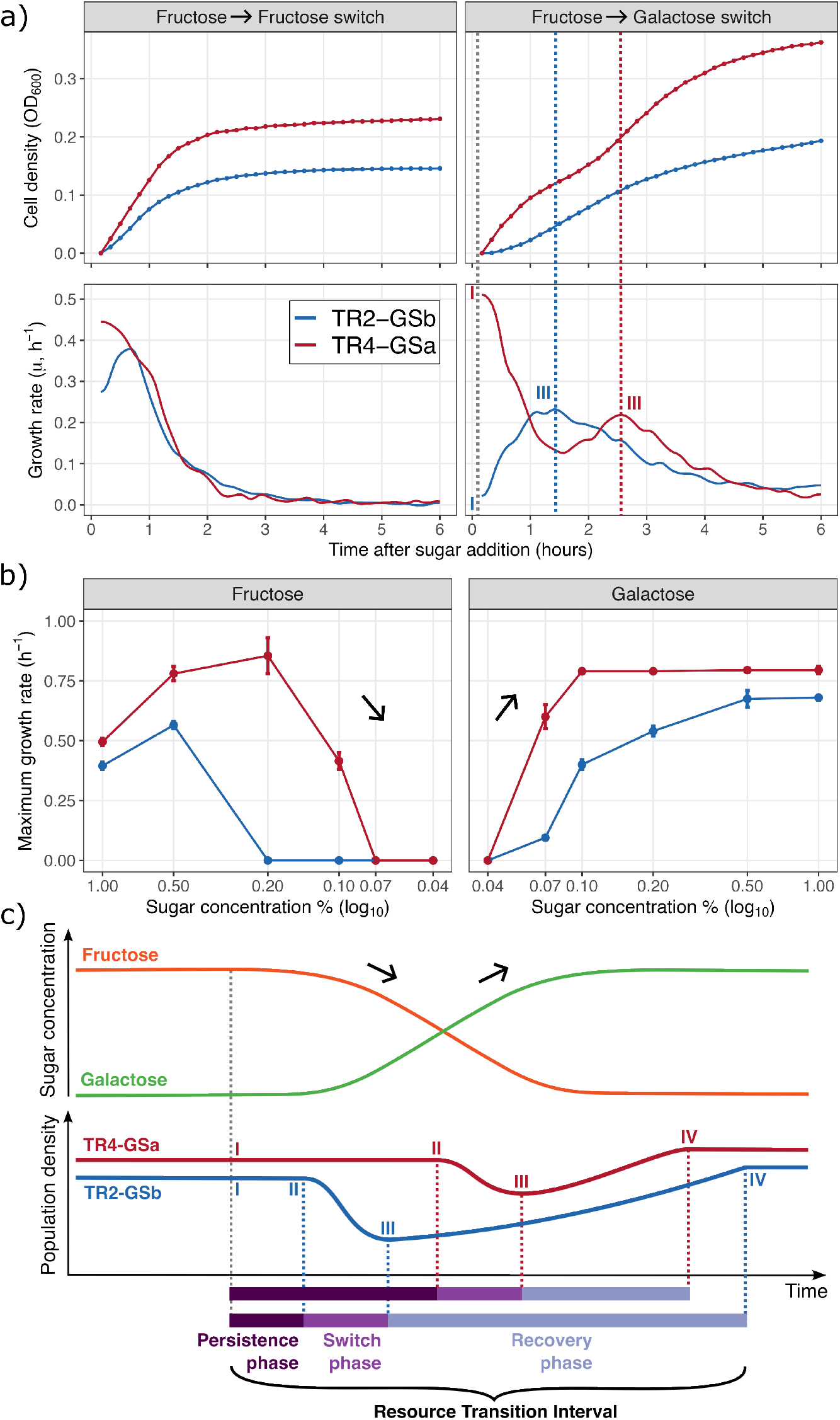
Differences in metabolic switching from fructose to galactose between two strains: TR2 (blue lines) and TR4 (red lines) sampled at the final timepoint (T3). **a)** Top panels: cell density during resource transition, calculated as cumulative growth yield normalized by starting optical density (OD). Bottom panels: specific growth rate, *µ* = d(ln OD_600_)*/*d*t*, estimated for each well by fitting a smoothing spline to ln(OD_600_) (not normalized) as a function of time and evaluating its first derivative. Dotted vertical lines indicate the beginning (I) and completion (III) of the resource transition based on achieving the maximum growth rate on the new resource. (see panel C). Left panels: no sugar switch (remains on fructose). Right panels: sugar switch from fructose to galactose. **b)** Evolved reaction norms inferred from resource-affinity measurements across gradients of fructose (left panel) and galactose (right panel) concentrations. Note that concentration ranges for the fructose panel (left) are reversed (decreasing) in order to visually match the directionality of a resource transition in the chemostat and panel C. Black arrows indicate if resource is increasing or decreasing. **c)** Conceptual model of resource and population dynamics in the chemostat during the transition from fructose to galactose. Fructose represented by the orange line, and galactose by the green line. Dotted vertical lines delimit the three phases (persistence, switch, and recovery) of the resource transition interval. Strain labels follow this scheme: the first letter indicates the treatment (T = temporal), followed by the replicate number (R1-R4). FS or GS after the dash denotes fructose or galactose specialists, respectively, and a final lowercase letter identifies the strain.

### 3.7 A conceptual model of resource transitioning in the chemostat

We propose a conceptual model in which the resource transition interval (i.e., the amount of time it takes for a population to regain steady state growth after a resource transition) is subdivided into three phases (Fig. 7c): 1) a persistence phase (current resource is decreasing in abundance but sufficient growth can still be maintained, 2) a switch phase: the population initiates a switch in expression towards an alternative resource strategy and 3) recovery phase: after completion of the resource switch the population regains balanced steady state between population and resource abundance. These three phases are marked by four events: onset of environmental resource transition (I), start of metabolic switch (II), completion of resource switch (III), and regaining of steady state between population and resource abundance (IV). The total length of the resource transition interval depends on two factors: the level of resource persistence (i.e., the ability to postpone a resource switch by continuing to grow at diminishing concentrations of resources) and the speed of resource plasticity (switching speed). This involves the temporal interaction between resource utilization efficiency of both fructose and galactose. The model reflects the dynamics of resource transitions in the chemostat. As the sugar input switches from fructose to galactose via the drip-feed mechanism, fructose levels drop rapidly through a combination of dilution and consumption, while galactose concentrations increase slowly as fresh medium enters. In this scenario, fructose levels are expected to decline faster than galactose levels rise because of ongoing consumption. Strains forced to switch early to galactose due to low resource utilization efficiency for fructose (e.g., TR2-GSb) may become temporarily trapped under conditions of insufficient galactose availability. Thus, if the resource utilization efficiency for galactose does not sufficiently match the encountered build-up concentration at the switch, the strain will be unable to sustain high growth rates until galactose accumulates. As a result, population density declines until galactose concentrations reach levels sufficient to restore growth and eventually steady-state resource and population abundance. Thus, evolving and maintaining a reaction norm that supports sufficient growth rates at lower resource concentrations provides a clear selective advantage through reducing variance of fitness by shortening the resource transition interval and minimizing the dip in growth (Fig. 7c). Such adaptations are expected only when a resource transition results in a lag in growth. If resource transitions do not lead to a decrease in growth rate, as was observed for the galactose-fructose transition in the ancestor strain (Supp. Fig. S7, S8), such adaptations are not expected to be selected for.

## 4. Discussion

### 4.1 Temporal variation selects variance-minimizing adaptations by equalizing growth performance across resources

Evolutionary theory predicts that in fluctuating environments, long-term fitness depends on the geometric mean performance across states, favoring strategies that reduce variance in success without necessarily maximizing arithmetic mean performance (Bradshaw, 1965; Gillespie, 1973; Bulmer, 1994; Kassen, 2002). Such a reduction in fitness variance can arise via fixed generalism, phenotypic plasticity (with switching costs), or bet-hedging. Consistent with these expectations, temporal fluctuations in fructose and galactose availability drove the repeated emergence of galactose specialists (GS), whereas the constant mix treatment maintained a stable polymorphism of GS and fructose specialists (FS). Given that the ancestor exhibited faster growth on fructose (0.82 h^−1^) than on galactose (0.43 h^−1^), selection under temporal variation favored elevated galactose performance, effectively equalizing growth across resources and thereby lowering fitness variance (Fig. 3a,d). By contrast, the mix treatment did not evolve performance equalization (Fig. 3b). This outcome aligns with the theory that, under constant co-availability and strong trade-offs, negative frequency dependence will tend to sustain a polymorphism of complementary specialists rather than a single phenotype that compromises on a generalist strategy. Additionally, the slower emergence of the GS phenotype in the mixed treatment could be explained by demographic factors: a smaller effective population size of the GS phenotype reduces the rate of adaptation (Fig. 3b). Together, these results demonstrate how temporal variation drives evolution toward variance-minimizing solutions through performance rebalancing. In contrast, constant environments with simultaneous resources allow for the coexistence of specialists, shaped by trade-offs.

### 4.2 Constraints on phenotypic plasticity depend on the directionality of switching

We observed a consistent asymmetry in switching behaviour under temporal variation: transitions from galactose to fructose occur without detectable decrease in growth rate, whereas those from fructose to galactose resulted in a clear growth lag (Supp. Fig. S8). Switching between resources that require distinct import systems and enzymatic conversions demands transcriptional investment of the cell in new metabolic machinery. The greater the number of differential metabolic steps, the higher the energetic cost and, thus, the more pronounced the growth lag. Because fructose metabolism involves relatively few metabolic steps compared to galactose (given the metabolic network structure of *L. cremoris*; Supp. Fig. S1; Ekkers et al. 2022), it is plausible that switching from galactose to fructose demands lower anabolic investment than the reverse, explaining the differences we observed in the growth lag under temporal resource variation.

These findings highlight that the cost of switching between phenotypic states is direction-dependent, a property that is generally not incorporated in classical reaction norm theory (Dupont et al., 2024). Standard models typically treat plasticity as a reversible, symmetric process, where a phenotype can be adjusted with equal facility regardless of the direction of change. However, Dupont et al. (2024) argues that such static representations overlook the temporal dynamics of plastic responses, in which both the speed and reversibility of switching are constrained by underlying physiological mechanisms. In our case, the asymmetric lag between the transitions galactose-to-fructose and fructose-to-galactose illustrates precisely this point: plasticity is not simply a matter of reaching a new phenotypic state, but of traversing a metabolic trajectory whose energetic and transcriptional demands depend on the direction of change. This directional dependence implies that fitness costs of switching are unequally distributed between alternative phenotypes: a galactose specialist may incur lower penalties when resources fluctuate, whereas a fructose specialist experiences stronger lags that reduce competitive performance. Similar switching asymmetries have also been documented in higher organisms. In birds, for example, seasonal switches between insects and fruits are easier in one direction than the other, because of beak shape constraints on handling times and diet breadth (Carnicer et al., 2009). Such asymmetries in switching costs may constitute a key trait component shaping the relative evolutionary success of competing phenotypes under temporally variable environments.

### 4.3 The evolution of reaction norms is accompanied by trade-offs across the environmental gradient

Rather than a uniform up- or downward shift in performance, we observed concentration-specific gains and losses in maximum growth rate: most strains (6 out of 8) showed their largest growth improvement at ≤ 0.5% resource concentrations, and most trade-offs (4 out of 7) were strongest in this same low-concentration range. This pattern is consistent with a combination of allocation, acquisition, and specialist–generalist trade-offs (Angilletta et al., 2003): selection in chemostats likely targeted the steady-state resource window, favoring genotypes that reallocate transcriptional and energetic budgets and/or enhance uptake at low concentrations, while sacrificing performance at higher concentrations. A striking example of such a trade-off was described for E. coli adapted to lactose-limited continuous culture, where subsequent exposure to high lactose concentrations caused severe growth inhibition and mortality (“lactose killing”) due to excessive sugar transport through lactose permease (Dykhuizen and Hartl, 1978). This illustrates how adaptation of resource acquisition to low substrate concentrations can become detrimental at high resource availability. A similar mechanism could potentially contribute to the reduced growth observed at high resource concentrations in some of our evolved strains, although we did not measure sugar uptake or cell viability and therefore cannot establish whether the underlying mechanism is comparable.

Notably, different proximate mechanisms can produce superficially similar reaction norms (Angilletta et al., 2003). In our case, the observed trade-offs within-resource utilisation reaction norms (Fig. 6; Supp. Fig S4) likely reflect distinct mechanistic routes, such as transporter affinity versus capacity, catabolite repression thresholds, or enzyme expression costs. While these routes produce comparable gains under the induced selection conditions (Fig. 3), they show divergent outcomes in the other phenotypic analyses (Fig. 4, 5, 7). Regarding functional traits, these concentration-dependent improvements suggest underlying mechanistic trade-offs operating across the resource gradient. These concentration-specific trade-offs could be related to regulatory shifts in sugar-import systems: up- or down-regulating transporters with different affinities can enhance performance in one concentration range while reducing it elsewhere (Montaño-Gutierrez et al., 2022; Bosdriesz et al., 2018; Jeckelmann and Erni, 2020). Indeed, it is known that *L. cremoris* features different import systems for fructose (Benthin et al., 1993) and galactose (Solopova et al., 2018; Neves et al., 2010). Moreover, our framework demonstrates that single-resource concentration or maximum growth rate assays - common in evolutionary studies - can overlook dominant phenotypic changes that occur at low concentrations (≤0.5%; Fig. 6). Incorporating full concentration gradients into assays thus provides a more accurate assessment of the trade-offs that shape reaction norms under realistic ecological constraints, aligning with Angilletta et al. (2003), who advocated for a unified, trade-off–centric view of reaction norm evolution.

### 4.4 Adaptation of growth persistence relaxes the need for speed

Temporal fluctuation favoured performance equalisation across fructose and galactose (Fig. 3a), but lineages reached this outcome via distinct routes (Fig. 6): the GS lineage TR2 switched early at the cost of a lower yield, whereas the generalist-like TR4 switched later, maintaining growth at lower resource concentrations. In the framework of reversible phenotypic plasticity (RPP) rates, performance in fluctuating environments depends not only on the extent of plasticity a trait can express (i.e., magnitude), but also on how quickly it can be expressed and the associated costs of that speed (Burton et al., 2022). TR2 resembles a high-rate, high-cost solution, whereas TR4 increases persistence as resources decline, buffering the phenotype-environment mismatch so that a slower switch still maximises geometric-mean fitness across resource fluctuations. This mechanism bears similarity with plasticity-mediated persistence (PMP), whereby plastic traits sustain individuals through adverse phases, minimising selection bottlenecks and providing time for adaptive tracking to occur (Morris, 2014). PMP, RPP, and our persistence-versus-switching dynamics thus share a common function: both buy time and broaden the tolerable mismatch between phenotype and environment.

Real-world resource transitions typically follow a pattern of decline and build-up. In seasonal systems, for example, fruit availability increases gradually, making switching too early costly (i.e, incoming resource has not yet reached concentrations sufficient to support performance), while switching too late risks missing the peak (De Villemereuil et al., 2020). Birds with traits that sustain performance on the declining resource (e.g., maintaining insect foraging efficiency as fruits begin to appear) can delay switching until fruits are abundant, thereby maximizing geometric-mean fitness across seasons (Carnicer et al., 2009). Our findings support similar dynamics, where persistence can substitute for speed: delayed switching remains optimal as long as it buffers phenotype-environment mismatch (Fig. 6). By prolonging effective use of the depleting resource, persistence reduces time spent in dual suboptimal states (too little of the old resource and not yet enough of the new), prevents premature switching, and allows the new resource to accumulate before a plastic switch is triggered. In RPP terms, enhanced persistence reduces the penalty of lag and increases the perceived predictability of the environment (Burton et al., 2022), offering a general alternative to evolving uniformly faster switching. More broadly, under climate-driven shifts in resource-availability windows, selection may favour greater persistence at low availability rather than uniformly faster plasticity - especially when rapid RPP is costly or physiologically constrained (Burton et al., 2022). Such persistence-mediated buffering is a plausible, general route to PMP in changing environments (Morris, 2014) and cautions against evaluating adaptive potential solely by maximal performance or instantaneous switching speed.

## 5. Conclusion

In our evolution experiment, selection for growth under temporal variation in resource supply promoted variance-minimization by equalizing performance across resources, consistent with a maximization of geometric mean fitness. Lineages displayed concentration-specific adaptations to the reaction norms for resource utilisation, which coincided with trade-offs at other concentration ranges. Our results show that temporal variation imposes multi-dimensional selection pressures, extending beyond a simple optimisation of maximal growth rates - it exposes populations to dynamic conditions, selecting also for growth yield, switching kinetics, and resource persistence. Trade-offs between these different fitness components allow for a diversity of qualitatively different resource-utilization strategies to thrive in a temporally variable environment, as reflected by the diversity of strategies observed in our experiment.

## Data availability

The data are not publicly available online but can be shared upon request by contacting the corresponding author.

## Acknowledgements

We thank Cyrus Mallon for help with running the evolution experiment, and Anne-Marie Veenstra-Skirl for assisting with the plating experiments. This work was supported by the European Research Council (ERC Starting Independent Researcher Grant 309555 to G.S.v.D.) and the Netherlands Organization for Scientific Research (NWO Vidi Grant 864.11.012 to G.S.v.D.).

## Author contributions

D.M.E. conceived and designed the study, performed the experiments, phenotypic characterization, led the data analyses, and wrote the manuscript. M.C.R. analysed the data (reaction norms), performed the clustering analysis, co-created the figures, and edited the manuscript. S.M.-G. analysed the data (growth curves, growth recovery, and regression analyses of the pH data). O.P.K. provided laboratory support, and experimental and genetic expertise for the conception of the project. G.S.v.D. co-conceived the idea, provided advice throughout the experiment, and edited the manuscript. All authors contributed to the interpretation of the data and commented on the manuscript.

