## Supplementary Information for "Temporal resource variation promotes growth equalization through alternative plasticity strategies"

### Supplementary Material

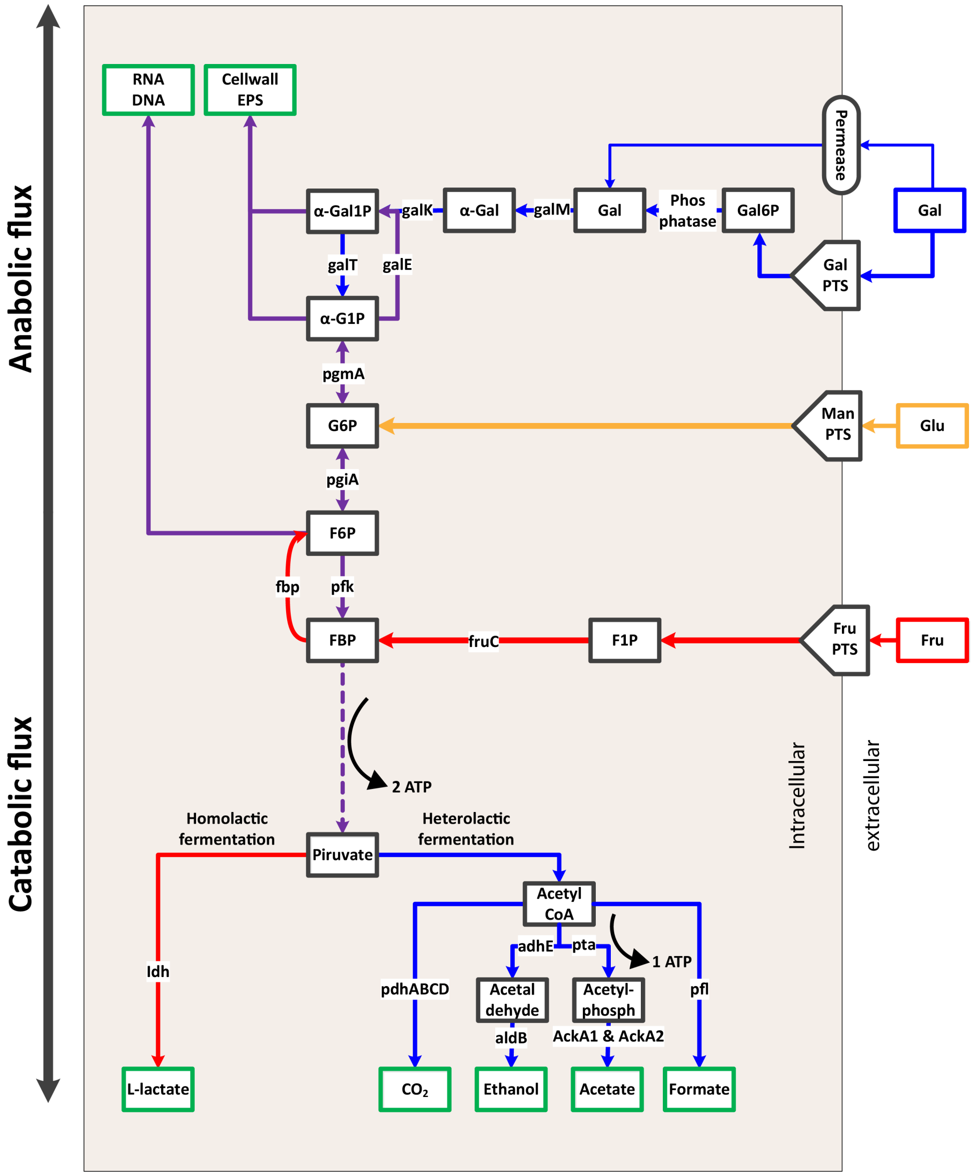


**Figure S1 Schematic representation of the metabolic architecture of the central carbon metabolism of L. cremoris MG1363.** The metabolic fluxes are visualized by color-coded arrows indicating the net direction of metabolic flux of galactose (Gal; blue), fructose (Fru; red) and glucose (Glu; orange - the sugar that the ancestor is preadapted to). Shared pathways are shown in purple. Text on the arrows indicates gene names of the enzymes related to each metabolic step. PTS: phosphotransferase system; EPS: extracellular polymeric substances. Metabolites are shown as squares: Gal6P: galactose-6-phosphate; Gal: galactose; $\alpha$-Gal: alpha-galactose; Gal1P: galactose-1-phosphate; G1P: glucose-1-phosphate; G6P: glucose-6-phosphate; F6P: fructose-6-phosphate; F1P: fructose-1-phosphate; FBP: fructose-1-6-biphosphate. Galactose enters glycolysis upstream of the anabolic junctions, whereas fructose enters downstream, requiring fluxes in upper glycolysis to be adjusted in opposite directions to maintain an optimal balance between anabolic and catabolic fluxes. Moreover, fructose and galactose metabolism also differ in terms of their anabolic investment in the pathways for processing each sugar. Fructose requires eight enzymes for its catalysis up until pyruvate, while galactose requires 14 steps. Due to this difference in enzymatic investment, metabolic transitions towards fructose utilization are expected to be faster than those towards galactose metabolism. See (Ekkers et al., 2020) for more details.

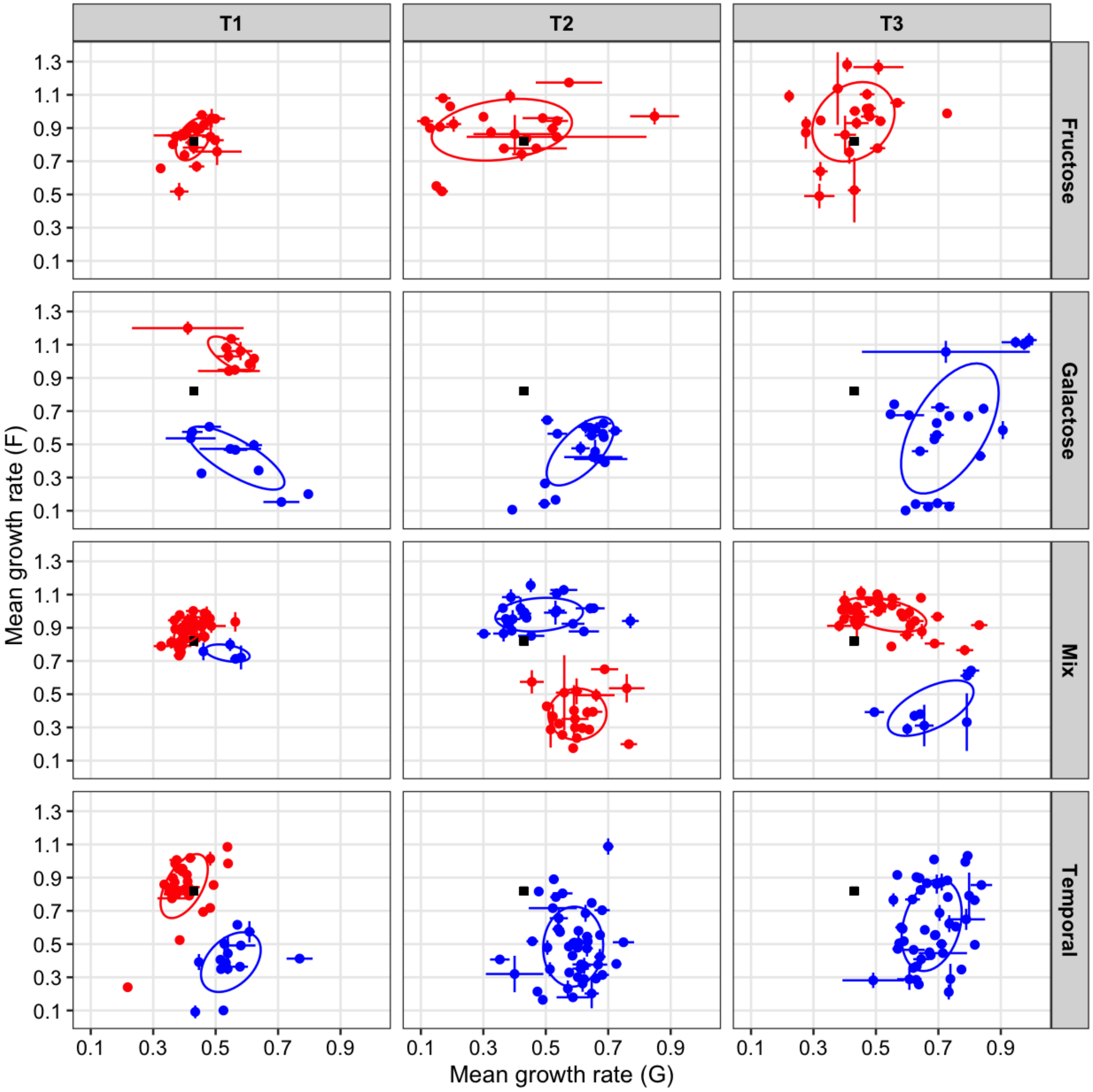


**Figure S2 Phenotypic clustering analysis.** Clustering was performed on the maximum growth rate coordinates on fructose (y-axis) and galactose (x-axis) for each of the genotypes sampled from the evolving populations (Fig. 2e). T1, T2, and T3 indicate the three sampled timepoints.

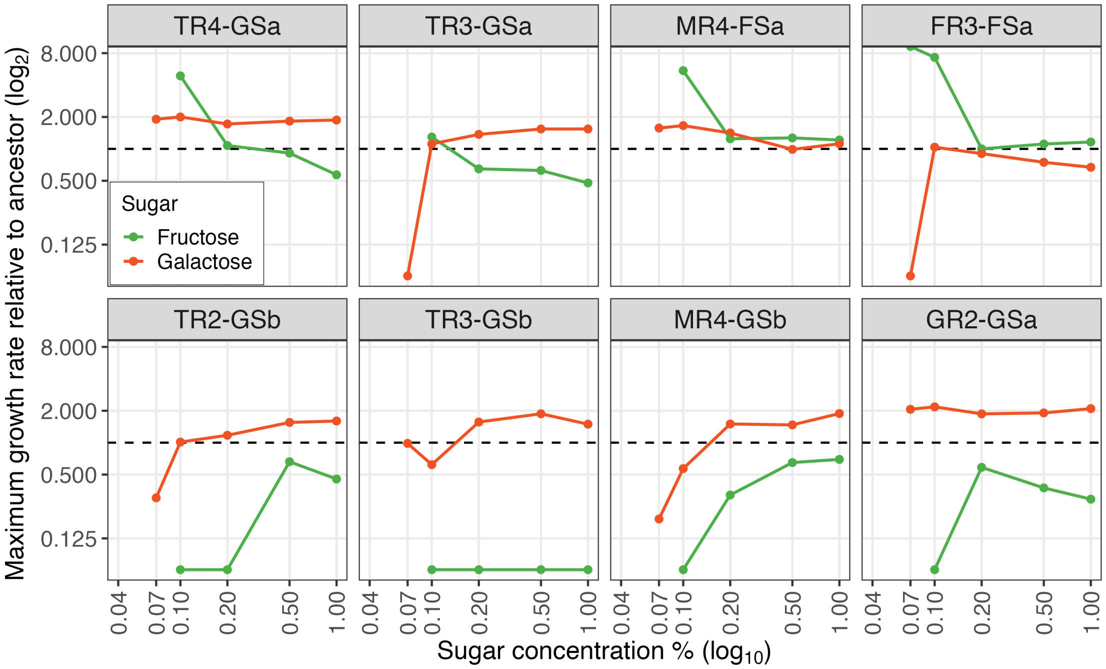


**Figure S3 Changes in resource utilization efficiency compared to the ancestor (ratio)**. The maximum rate of growth was measured at a gradient of resource concentrations (% w/v) of the ancestor and eight evolved strains in CDMPC. At each concentration, growth rates of the evolved strains were divided by ancestor growth rate to indicate relative change along the resource gradient. Growth rates are in $\log_2$. Strain labels follow this scheme: the first letter indicates the treatment (M = mix; F = only fructose; G = only galactose; T = temporal), followed by the replicate number (R1-R4). FS or GS after the dash denotes fructose or galactose specialists, respectively, and a final lowercase letter identifies the strain. Strains were sampled at the final timepoint (T3). No growth was measured in all strains at 0.04% sugar. See also Fig. 6.

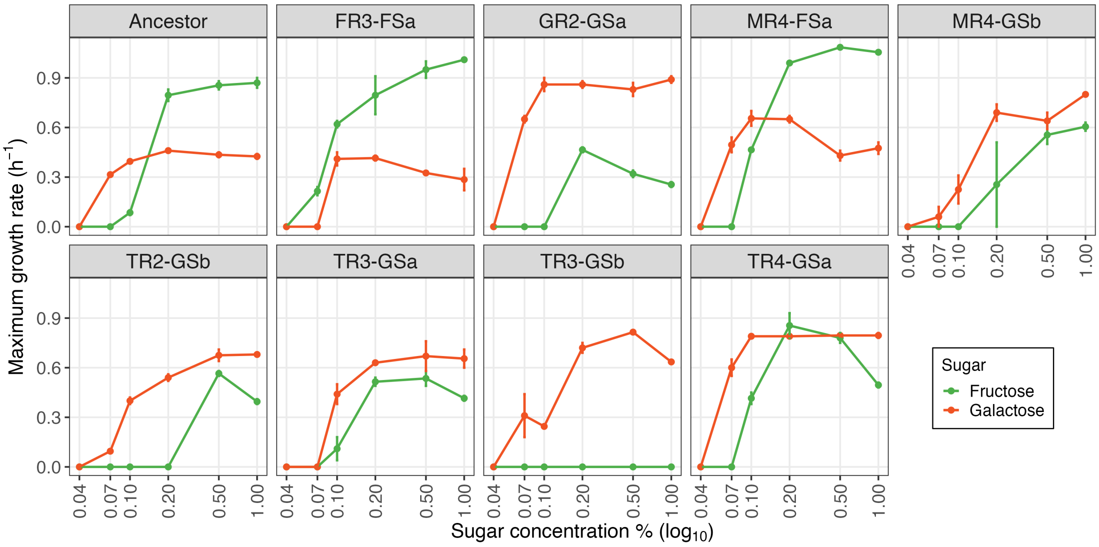


**Figure S4 Reaction norms of resource utilization efficiency**. Maximum rate of growth at a gradient of resource concentrations (% w/v) of the ancestor and eight evolved strains in CDMPC. Strains were sampled at the final timepoint (T3), and strain labels follow the scheme of Sup. Fig. S3. See also Fig. 6.

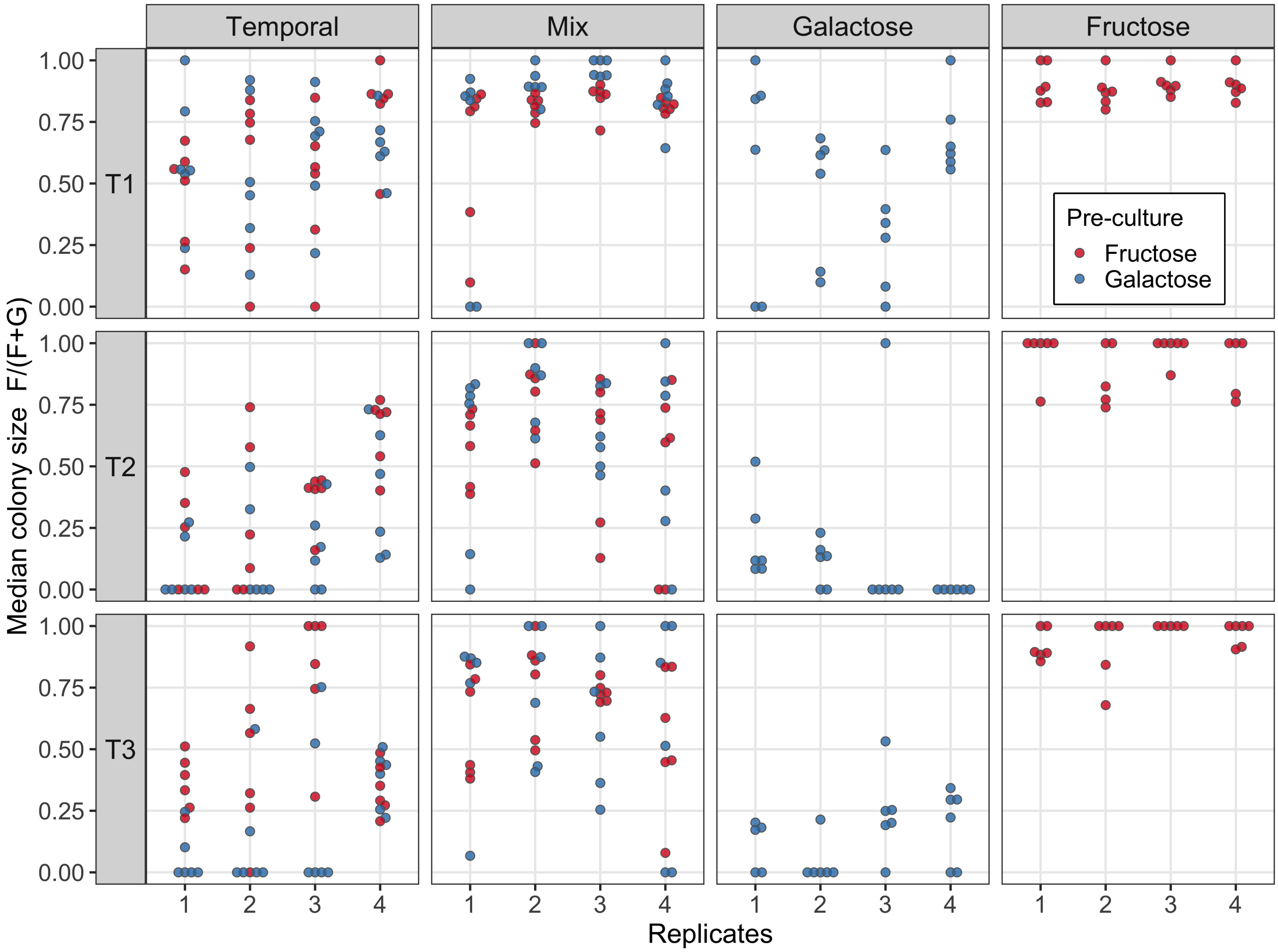


**Figure S5 Degree of specialisation to fructose based on differences in median colony size of evolved strains growing in fructose- versus galactose-supplemented medium**. Median colony size on fructose was divided by total median size (i.e., on fructose plus on galactose) to create a ratio that indicates relative colony size between both resources (1: only able to grow on fructose; 0: only able to grow on galactose; 0.5: equal colony sizes on both sugars). T1, T2 and T3 indicate the three sampled timepoints.

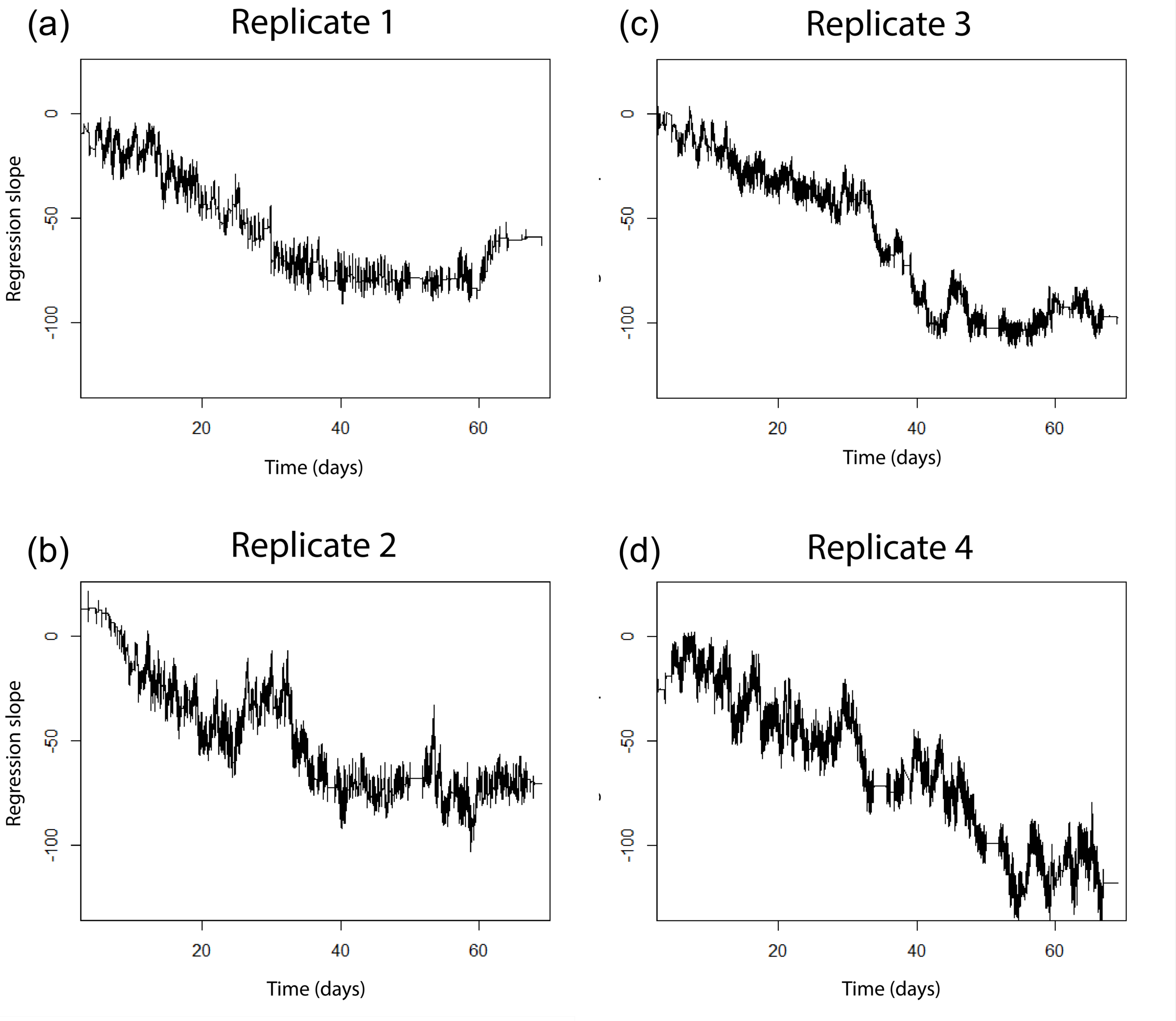


**Figure S6 Change in acidification rate throughout the evolution experiment.** The acidification rate of individual chemostats followed a pattern of steady increase of acidification throughout most of the evolution experiments ending with a gradual decrease in the change of acidification towards the end. Replicate 4 stood out because the rate of change acidification progressed longer and and reached a higher rate of acidification compared to the other replicates. Acidification rate was estimated by fitting a linear least-squares regression to each interval measurement interval. Precise quantification of the acidification rates was impeded by sensor noise.

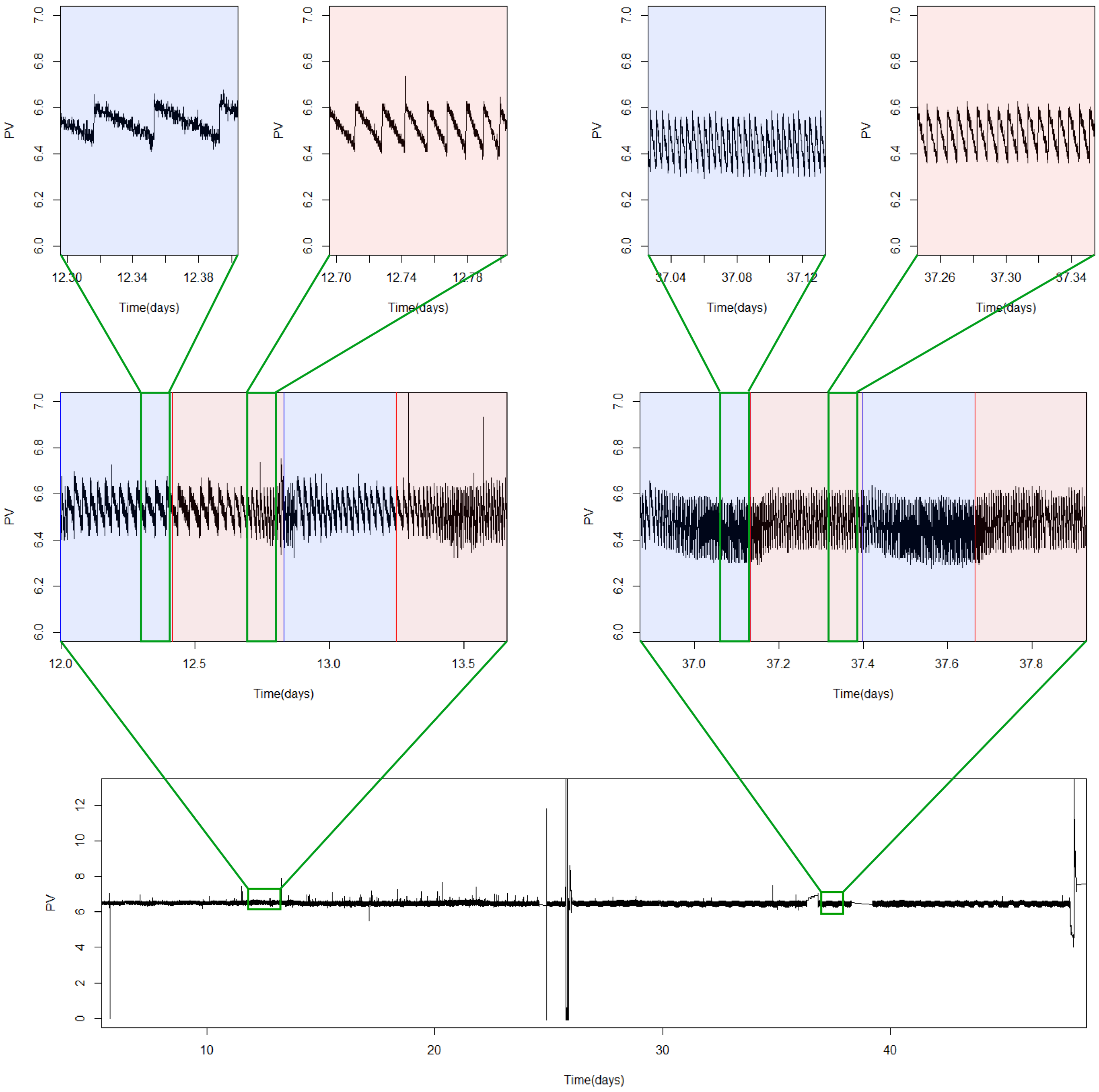


**Figure S7 Acidification rate of chemostat cultures.** Metabolic acidification patterns during fructose and galactose metabolism. pH sensor data are from the temporal chemostat treatment. Background shading indicates which sugar was supplied. The pH was maintained at 6.5 by a peristaltic pump that added NaOH in response to acidification (via PID loop) caused by metabolic waste production. The sensor data show a repeating pattern of gradual pH decline followed by a sharp increase. The gradual decrease reflects the accumulation of acidic metabolites, while the sudden increase marks the moment a droplet of NaOH is added to the culture. Both the slope of acidification and the frequency of NaOH additions serve as proxies for growth rate, reflecting the link between biomass production and waste metabolite accumulation. Measurements were taken at the end of each cycle to avoid interference from metabolic shifts. **(a)** Comparing the fructose and galactose supply phases, *L. cremonis* initially has a 2.7 fold higher acidification rate on fructose (pink shading on background) than on galactose (blue). **(b)** However, this growth asymmetry reverses towards a 1.8 fold higher acidification rate during growth on galactose in the second half of the experiment in all (G : F ratio 1.8 : 1), indicating that the evolving populations improve their performance on galactose relatively more than on fructose.

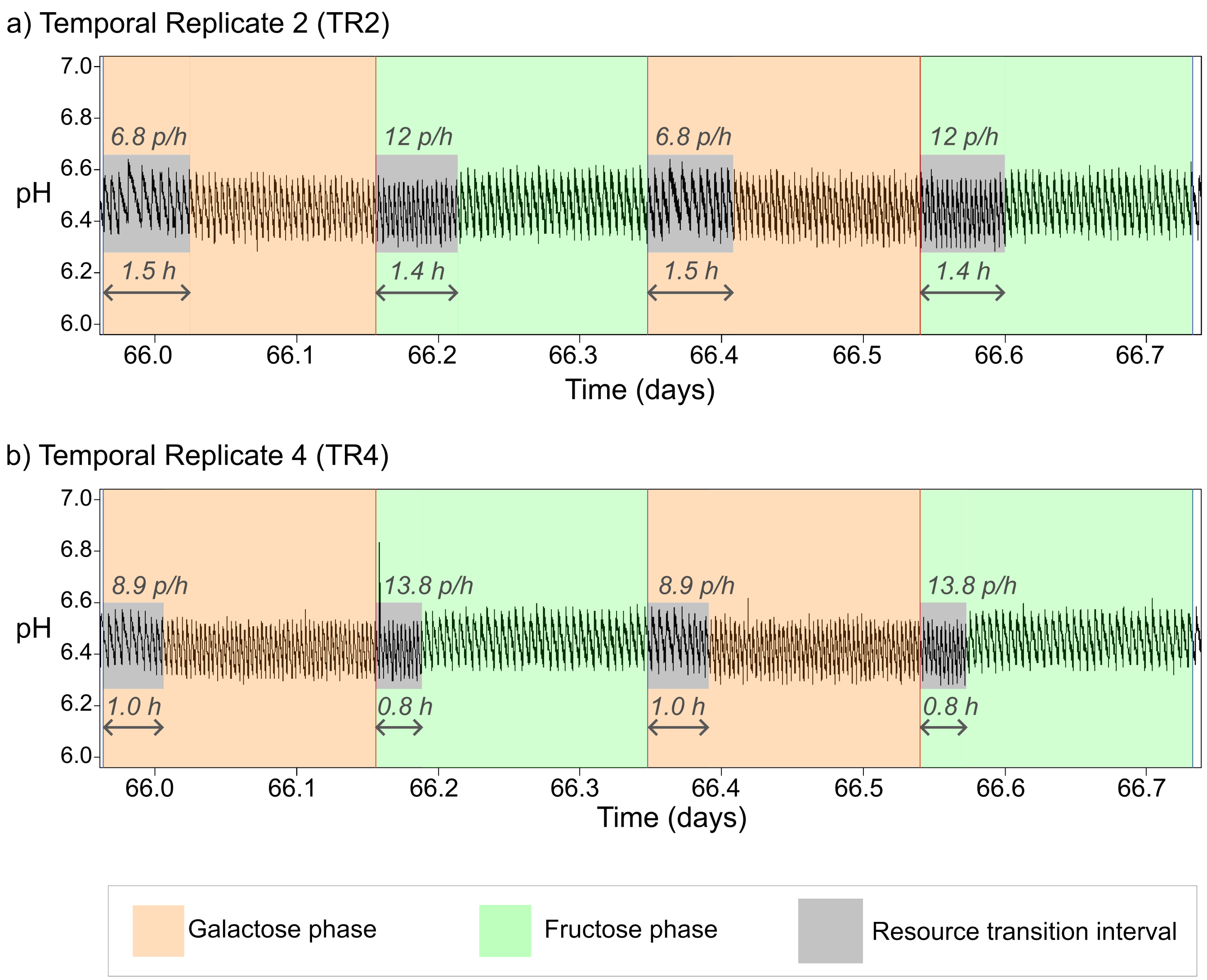


**Figure S8 Acidification rate during the resource transition interval.** Plasticity in metabolic switching between the fructose and galactose resources during the temporal treatment during the evolution experiment. Two replicate chemostats (TR2 and TR4) are compared towards the end of the experiment (around timepoint T3) in terms of their resource transition length and the acidification rate per unit of time within this interval. The resource transition interval (in grey) starts at the moment the resource valve is activated to switch the resource (i.e., beginning of each coloured phase: orange for galactose, green for fructose). The end of the resource transition interval is defined by the moment that the culture re-established a steady-state acidification rate, determined by eye. The steady-state growth interval is defined by the time between the start of steady state growth until the next resource switch. The acidification rate is calculated by calculating the amount of NaOH droplets (inferred from pH sensor data) per unit of time (unit: peaks of pH per hours; p/h). We find an asymmetry between switch direction for both chemostats, in which switching from galactose to fructose appears to occur without a reduction in metabolic activity, whereas switching from fructose to galactose results in a notable decrease in acidification rate, before establishing a steady state growth rate. In general we find that TR2 shows a lower acidification rate and longer resource transition interval compared to TR4, this is particularly visible for the transition from fructose to galactose.
